# Divergent co-transmission by a predictive motor circuit modulates auditory processing

**DOI:** 10.64898/2026.01.22.700061

**Authors:** Marryn M. Bennett, Han S.J. Cheong, Kaitlyn N. Boone, Farzaan Salman, Jacob D. Ralston, Jenna M. Luthman, Kevin C. Daly, Gwyneth M. Card, Andrew M. Dacks

## Abstract

During behavior, an animals’ movements often stimulate its own sensory receptors. For instance, whether during flight or the production of courtship song, when flies beat their wings they cause mechanosensory disturbances that activate primary auditory neurons in the antennae. However, flight and courtship occur under very different behavioral contexts and temporal-scales. As a consequence, the nervous system contains predictive motor circuits that modify sensory processing in anticipation of self-induced (re-afferent) stimulation. However, it is unclear how information about planned movements can be used to effectively tune specialized sensory processing circuits. Here we describe a predictive motor circuit that releases two transmitters to differentially modify auditory neurons that process sound under different behavioral contexts. In *Drosophila*, two metathoracic ascending histamine neurons (MtAHNs) become active prior to wing movement and project to the primary auditory network. We found that the MtAHNs co-transmit the neuropeptide Dh44 and histamine, and each suppresses the activity of auditory neurons that are integrated into distinct networks that support escape and courtship, respectively. Additionally, Dh44 has a greater impact on male auditory neurons and suppression by Dh44 occurs over a longer time course than histamine. Disrupting either histamine synthesis or neuropeptide release by the MtAHNs reduces courtship success with an increase in cessation during the wing extension step. Consistent with a role in auditory processing of courtship cues, Dh44 cotransmission is present in *Drosophila* species that have an auditory component to their courtship displays, but is absent in species that use either purely visual courtship or lack apparent courtship behavior. Finally, the projections of the MtAHNs into auditory centers were missing in snow flies, which are wingless. Our results indicate that divergent cotransmission by predictive motor circuits may represent an efficient means to differentially modulate sensory subnetworks impacted by reafferent feedback.

## Introduction

As animals navigate their environment, they must differentiate between sensory input that derives from the environment (exafference) and that which arises as a consequence of their own movement (reafference). The nervous system can execute this comparison through predictive signals, sent from motor networks to sensory circuits, called corollary discharge (Crapse & Sommer, 2008). Corollary discharge circuits (CDCs) impact sensory processing during a wide array of behaviors from eye saccades in primates(Sommer & Wurtz, 2002, 2004; Tanaka, 2006), motion processing in flies (Fenk et al., 2021; Kim et al., 2017) and stridulation in crickets(Poulet & Hedwig, 2006), and deficits in CDC function can be attributed to many forms of sensory hallucinations(Feinberg & Guazzelli, 1999; Ford & Mathalon, 2004; McGrath et al., 2015; Morris, 1960). Often CDCs rely upon a corollary discharge interneuron (CDI) to inhibit the sensory circuit that is immediately impacted by the ongoing behavior. For example, male crickets produce sound (“stridulation”) to attract females by rubbing their wings together. To avoid being deafened by their own call, the central pattern generator that coordinates the articulation of the wings during stridulation also activates a GABAergic CDI that inhibits the auditory afferents and interneurons that respond to stridulation produced by other males(Baden & Hedwig, 2010; Poulet & Hedwig, 2006). In this manner the cricket auditory system is gated by the GABAergic CDI for the duration of the behavior. However, reafference from one behavior may affect multiple sensory circuits that feed into separate, timescale-dependent behavioral pathways and it is unclear how CDCs can selectively influence each circuit.

In this study we describe co-transmission as a means for a CDI to separately and differentially impact downstream targets allowing for timescale dependent signaling through the use of fast ionotropic receptors or slow metabotropic receptors. Co-transmission can be convergent, downstream neurons express receptors for both transmitters, or divergent, separate downstream neurons express receptors for either transmitter (Svensson et al., 2019). For instance, motor neurons of larval *Drosophila melanogaster* (Luo et al., 2017; Ormerod et al., 2016) and cockroaches (Adams & O’Shea, 1983; Nässel, 2018) co-transmit glutamate and proctolin which produce short and long term muscle contractions within the same myocyte, respectively (Adams & O’Shea, 1983; Ormerod et al., 2016). In mammals wakefulness is promoted through the co-transmission of glutamate and hypocretin/orexin, which respectively cause short and long-term excitation of histamine neurons in the tuberomammillary nucleus(Huang et al., 2001; Schöne et al., 2014). Thus, convergent co-transmission allows for distinct, immediate and delayed effects on a single target. Conversely, divergent co-transmission allows for one neuron to affect separate networks on distinct timescales. For example, maintenance of the gastric mill rhythm (GMR) in the crab, *Cancer borealis,* is mediated by co-transmission of GABA and cabTRP la from the projection neuron MCN1. GABA provides fast excitation to the GMR generator INT1 through ionotropic receptors and cabTRP la provides delayed excitation to the GMR generator LG through metabotropic receptors(Coleman et al., 1995; Stein et al., 2007). Therefore, divergent co-transmission allows neurons to segregate their influence by impacting distinct downstream targets on separate timescales. Here we describe how divergent co-transmission is potentially used by a CDI to promote sensory modulation in two subdivisions of the auditory network in *Drosophila melanogaster*.

In this study we examine the role of divergent, inhibitory co-transmission within a CDC. We show that a CDI projecting from a wing motor control network in the ventral nerve cord to an auditory network in the brain co-transmits histamine and the neuropeptide Dh44. This CDI synapses upon neurons in the auditory system that participate in separate behavioral circuits and differentially express receptors against histamine and Dh44. We demonstrate that histamine and Dh44 alter sound responses of auditory neurons that contribute to distinct behavioral networks over distinct timescales. Finally, co-transmission of Dh44 correlates with the use of auditory communication in species specific courtship behavior. Thus, we demonstrate how divergent co-transmission is potentially utilized by a CDI to differentially impact the mechanosensory processing for distinct behavioral circuits.

## Results

We recently described a flight CDI consisting of two pairs of ascending histaminergic neurons (AHNs) in the ventral nerve cord (VNC) of *Drosophila melanogaster*(Cheong et al., 2024). The metathoracic AHNs (MtAHNs) become active just prior to the onset of flight and project to auditory processing regions of the brain. During flight, wing beating produces mechanical reafference at a frequency of 200Hz which falls within the response range of both subtypes of sound responsive Johnston’s organ neurons (JONs) in the antennae (Ishikawa et al., 2020; Iyengar & Wu, 2014; Kamikouchi et al., 2009; Mamiya & Dickinson, 2015). However, JON subtypes differentially contribute to the neural circuits responsible for distinct behaviors. For instance, the giant fiber neurons integrate mechanosensory input from vibration sensitive JONs and visual input from the optic lobes to drive escape behavior (Ache et al., 2019; Jezzini et al., 2018; Pézier et al., 2014; Von Reyn et al., 2017; Wyman et al., 1984). Likewise, courtship circuits also rely on input from the vibration sensitive JONs to auditory projection neurons (Baker et al., 2022; Kamikouchi et al., 2009; Matsuo et al., 2014; Tootoonian et al., 2012; Vaughan et al., 2014; Zhou et al., 2015a). Thus, mechanical reafference from flight has the potential to affect behavioral circuits that operate on different timescales and under different contexts. As potential CDIs that modulate sensory processing by both circuits, the MtAHNs may therefore need to provide distinct signaling to separate components within auditory networks. Therefore, we sought to determine if the MtAHNs are capable of implementing divergent co-transmission to target separate components of the auditory network.

### The MtAHNs co-transmit histamine and Dh44

The MtAHNs project throughout the VNC as well as within auditory sensory processing regions of the brain including the antennal mechanosensory and motor center (AMMC; Figure 1A). Inducing apoptosis by expressing “*reaper*” (Klemm et al., 2021; White et al., 1996) in the MtAHNs eliminated histamine labeling in the AMMC (Figure S1.4A-B). Thus, the MtAHNs are the sole source of histamine to the AMMC (Cheong et al., 2024). Within the AMMC, the top downstream targets of the MtAHNs identified by connectomic analyses are primarily sensory neurons, interneurons, and descending neurons (Figure 1B, Figure S1.3). If the MtAHNs release only histamine, then their downstream targets should all express one or both of the two known insect histamine receptors, *HisCl1* and *ort* (Zheng et al., 2002). However, the majority of the cell types identified as targets of the MtAHNs within the connectome were not expressed in reporter lines for *HisCl1* and *ort* within the brain (Figure 1C-D) or the VNC (Figure S1.1). Prior work has demonstrated that *ort* is expressed in photoreceptors (Gengs et al., 2002; Zheng et al., 2002) and transcriptomic analysis of the antennae indicated *HisCl1* is expressed by the JONs (Senthilan et al., 2012) which matched the expression of the *ort* and *HisCl1* reporter lines respectively (Figure 1D, Figure S1.2A-B). We therefore surmised that the MtAHNs express a second neurotransmitter.

**Figure 1.**
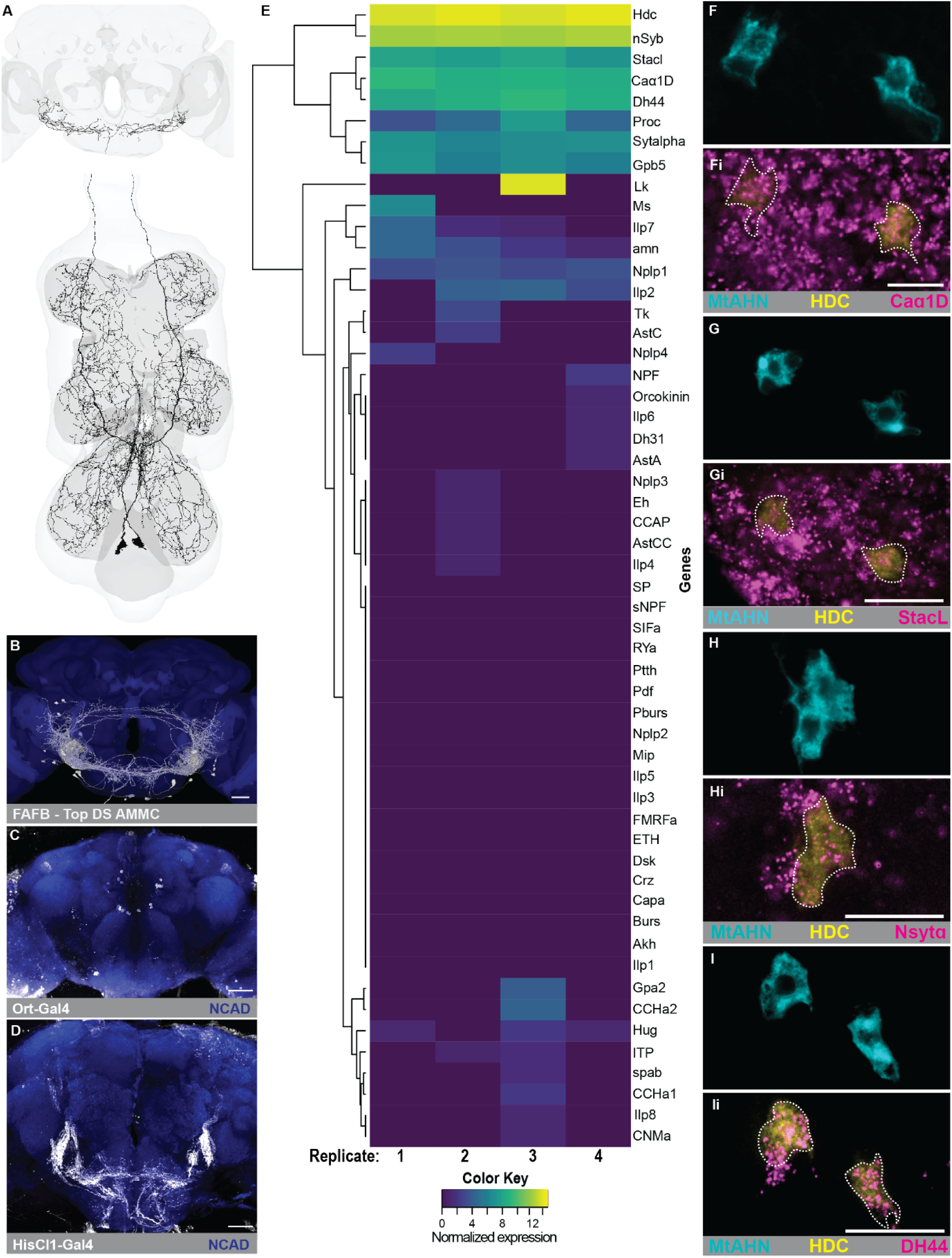
The MtAHNs co-transmit histamine and Dh44. **(A)** EM reconstruction of the MtAHNs in the brain (FAFB) and ventral nerve cord (FANC). **(B)** Top 9 cell types downstream of the MtAHNs in the AMMC (White) overlaid on neuropil mesh (Blue) from FAFB Flywire dataset. **(C)** TI{2A-Gal4}ort[2A-Gal4] driving expression of GFP (White) in the brain with NCAD (Blue) as a neuropil marker. **(D)** T2A-Gal4{HisClB}T2A-Gal4 driving expression of GFP (White) in the brain with NCAD (Blue) as a neuropil marker. **(B-D)** scale bars = 30μm. **(E)** Normalized expression from MtAHN RNAseq for neuropeptide and peptide release related genes with histidine decarboxylase (HDC) as a reference for histamine expression and Nsyb as a neuronal cell marker, log10 normalized expression level indicated with color key. **(F-I)** Single optical section images of GFP expression (Cyan) driven by VT025938AD ∩ VT040583DBD splitGal4 with HDC HCR-labeling (Yellow) as verification of histaminergic cells **(Fi)** Caα1D (Magenta), **(Gi)** StacL (Magenta), **(Hi)** Nsytα (Magenta), **(Ii)** Dh44 (Magenta) expression in the MtAHNs, cell body location indicated with white dotted line. **(F-Ii)** scale bars = 20μm.

Using a MtAHN driver line to express GFP, we isolated individual MtAHN somata for RNA sequencing. Using the neuronal cell marker nsyb and histidine decarboxylase (HDC; the rate limiting enzyme for histamine production) to verify cell identity, we queried against known transmitters and observed high expression of the neuropeptide *Dh44* and genes associated with the release of neuropeptide including *StacL* (Hsu et al., 2018, 2020), *Caɑ1D* (Hsu et al., 2020; Zupanc, 1996), and *Nsytɑ* (Park et al., 2014) across all replicates, as well as *leucokinin* in one replicate (Figure 1E). Antibody labeling for Dh44 does not overlap with GFP signal driven by a MtAHN driver line and neither does labeling for leucokinin (Figure S1.6D’-D’ii,E-Eii). However, histamine labeling in VNC of flies from a Gal4 reporter of endogenous Dh44 translation reveals overlap between histamine and Gal4 expression where the MtAHN cell bodies reside (Figure S1.6D”-D”ii). Thus, we reasoned that the Dh44 antibody may not effectively label cells with lower overall expression of Dh44. Therefore, we used a more sensitive labeling approach, hybridization chain reaction (HCR), to label *Dh44* and *HDC* mRNA. Dh44-Gal4 expression and *Dh44* HCR labeling in the VNC are highly overlapping (Figure S1.6D,D”), and there is overlap of HCR signal for *Dh44* and *HDC* mRNA transcripts in the MtAHN cells bodies (Figure 1I-Ii, Figure S1.6D-Dii). Consistent with the RNAseq data, the MtAHNs do not express additional small neurotransmitters (Figure S1.5) or neuropeptides (Figure S1.7). To confirm the capability of the MtAHNs to transmit neuropeptides we evaluated expression of several genes associated with the release of neuropeptides including the neuropeptide related synaptotagmins, Nsytalpha and Nsytbeta (Park et al., 2014), the peptidergic cell marker “DIMM”(Park et al., 2008), the L-Type calcium channel subunit *Caɑ1D* (Hsu et al., 2020; Zupanc, 1996), and the calcium channel regulator *StacL* (Hsu et al., 2018, 2020). Antibody and HCR labeling confirm that the MtAHNs express Nsytalpha but not Nsytbeta (Figure S1.6 A’-A’ii, C-Cii, Figure 1H-Hi, Figure S1.6A-Aii). HCR labeling for *StacL* and *CaAlpha1D* transcripts in VNC does overlap with GFP expression in the MtAHN cell bodies (Figure 1G-Gi, F-Fi), however a DIMM reporter line does not show overlap with the MtAHNs (based on histamine antibody labeling; Figure S1.6B-Bii). Overall, these results indicate that the MtAHNs are peptidergic and co-express histamine and Dh44.

### MtAHNs use co-transmission to modulate JO-As and JO-Bs on distinct timescales

The expression of histamine and Dh44 potentially allows the MtAHNs to differentially modulate their downstream targets. We therefore sought to evaluate this differential modulation within the context of a single sensorimotor circuit. Since connectomic analysis revealed that the MtAHNs are upstream of primary and secondary auditory neurons (Figure S1.3 A, B, F), we explored the impact of co-transmission by MtAHNs within the auditory network. The expression of HisCl1 by the JONs was previously established through transcriptomics (Senthilan et al., 2012) and we confirmed this through the HisCl1-Gal4 line (Figure 1D, Figure S1.2A-B). JONs are subdivided into subgroups based on their frequency tuning and AMMC projection morphology (Kamikouchi et al., 2006; Matsuo et al., 2014; Patella & Wilson, 2018), and make distinct contributions to different behavioral contexts (Baker et al., 2022; Kyriacou & Hall, 1982; Mamiya & Dickinson, 2015; Pézier et al., 2014; Zhou et al., 2015a). JO-As preferentially respond to frequencies ∼200Hz, JO-Bs respond to lower frequencies ∼100Hz, and JO-C through E respond to static deflections (Matsuo et al., 2014). We therefore first sought to determine which JON subtype expresses the HisCl1 receptor by using HCR to label mRNA transcripts of *HisCl1* in the second antennal segment (Figure 2A-C) of previously described JON-Gal4 lines (Figure S2.1) (Ishikawa et al., 2020; Kamikouchi et al., 2006) and calculating the proportion of HisCl1+ cells in each line. The proportion of expression of HisCl1 is significantly higher in the JO-A subtype when compared to the JO-B and JO-CE subtypes (Figure 2D), consistent with the observation from that the MtAHNs preferentially synapse upon the JO-A subtype in FAFB FlyWire(Cheong et al., 2024; Dorkenwald et al., 2022; Schlegel et al., 2024). We did not observe the expression of Ort in the JO (Figure S3.1E). Taking a similar approach to label for both Dh44 receptors (Cabrero et al., 2002; Hector et al., 2009; Johnson et al., 2005) in the JONs (Figure 3Ai-Ci, Aii-Cii) we observed that the JO-Bs have a significantly higher proportion of expression for both Dh44 receptors relative to the JO-As and JO-CEs (Figure 3E-F). Thus, the MtAHNs may use co-transmission to differentially target two separate JON subtypes, histamine for the JO-As and Dh44 for the JO-Bs.

**Figure 2.**
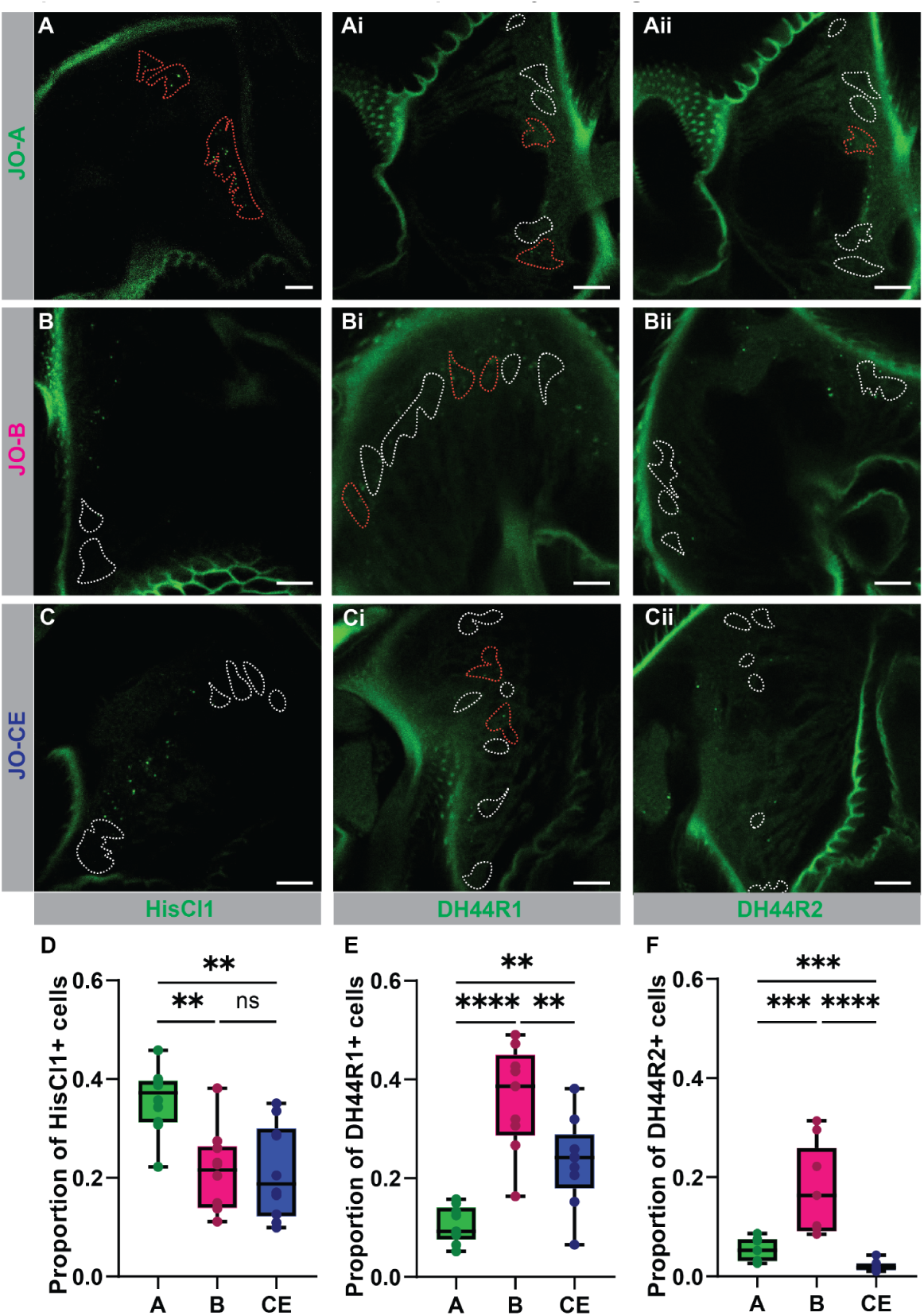
JONs types express different receptors for the transmitters of the MtAHNs. Cell body, single optical section, images of HCR labeling (green) for **(A-C)** HisCl1, **(Ai-Ci)** Dh44R1, **(Aii-Cii)** Dh44R2 in each JON cell type indicated on the left, positive cells indicated with red dotted lines and negative cells indicated with white dotted lines. **(D-F)** Box plots of proportion of expression in each JON subtype for **(D)** HisCl1 (N=10), **(E)** Dh44R1(N=9), and **(F)** Dh44R2(N=9) (significance from one-way ANOVA with post-hoc Tukey’s test for individual comparisons: NS=P>0.05, *=P<0.05, **=P<0.01, ***=P<0.001, ****=P<0.0001). Scale bars = 10µm.

**Figure 3.**
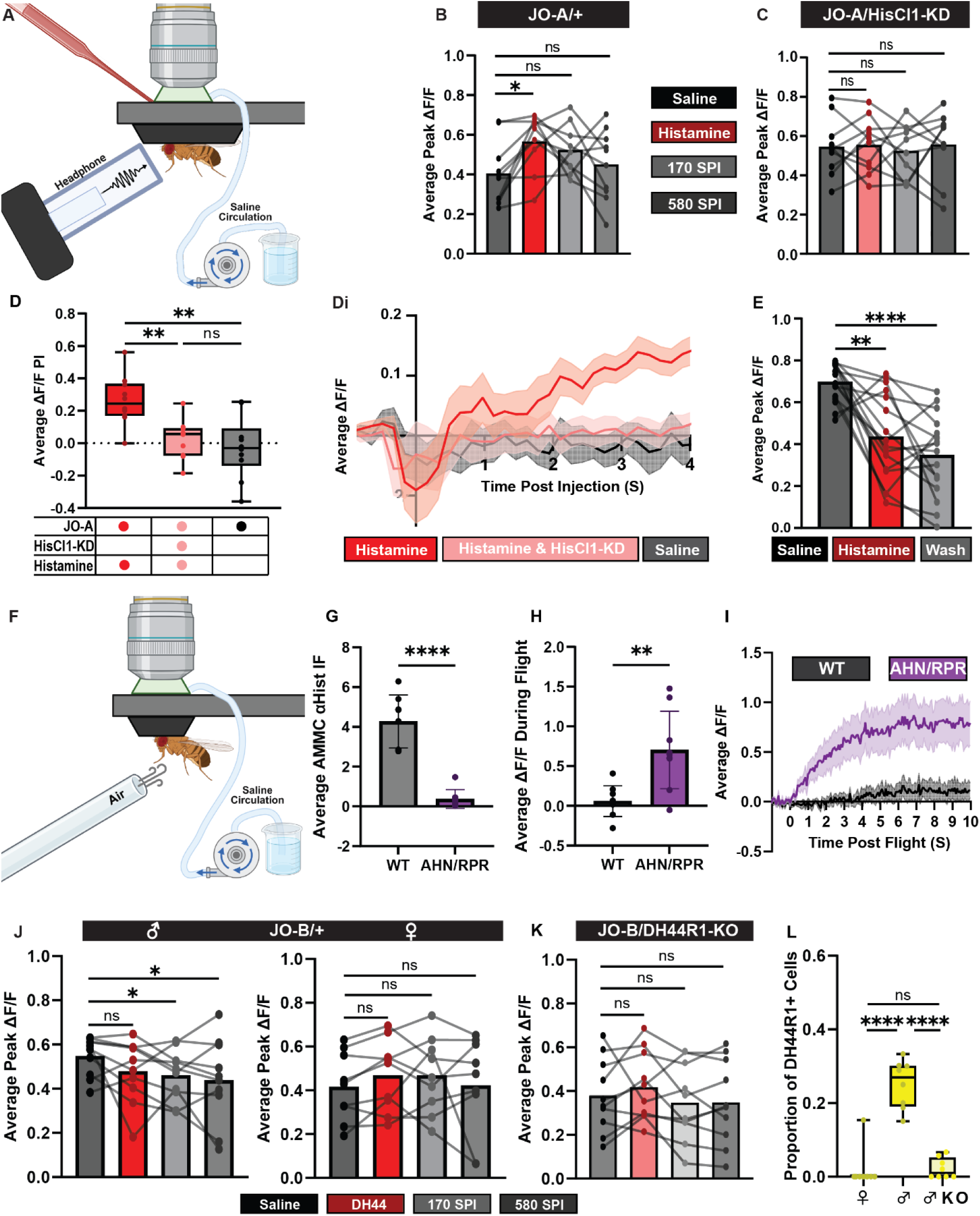
The JO-A neurons of both sexes exhibit PADs-like responses when exposed to histamine while the male JO-B neurons exhibit a reduction in sound response when Dh44 is applied. **(A)** Schematic of experimental set-up for Ca^2+^ recordings with sound presentation and pico injection. **(B-C)** Before and after plots of average ΔF/F during sound presentation in flies with **(B)** R74C10-Gal4 driving expression of GCaMP7f and **(C)** HisCl1-RNAi under saline conditions (black), pico injection of histamine (red), 170 seconds post injection (SPI) (light grey), and 580 SPI (dark grey)(N=10, significance from repeated measures ANOVA with post-hoc Tukey’s test for individual comparisons: NS=P>0.05, *=P<0.05). **(D)** Box Plots detailing the average ΔF/F post injection with the genetic and chemical variables indicated below (N=10, significance from one-way ANOVA with post-hoc Tukey’s test for individual comparisons: NS=P>0.05, **=P<0.01). **(Di)** Ca^2+^ traces (N=10, average ΔF/F ± SEM) prior to sound presentation from flies with R74C10-Gal4 driving the expression of GCaMP7f and pico injection of histamine (red), pico injection of histamine with HisCl1-RNAi (light red), and pico injection of saline (grey). **(E)** Before and after plot of average ΔF/F during sound presentation in flies with R74C10-Gal4 driving expression of GCaMP7f with initial saline application (black), 200µM histamine bath application (red), and saline wash (grey) (N=20, significance from repeated measures ANOVA with post-hoc Tukey’s test for individual comparisons: **=P<0.01, ****=P<0.0001). **(F)** Schematic of flight induction experimental set up. **(G)** Bar graph of average fluorescence intensity from histamine antibody labeling in the AMMC of wildtype (grey) and VT059420-LexAGADfl>LexAop-RPR (purple) flies (N=8,significance from unpaired t-test: ****=P<0.0001). **(H)** Average ΔF/F during flight in JO-A (R74C10>GCaMP7f) AMMC projections of wildtype (grey) and VT059420-LexAGADfl>LexAop-RPR (purple) flies with antenna glued (N=9,significance from unpaired t-test: **=P<0.01). **(I)** Ca^2+^ recording traces from JO-A (R74C10>GCaMP7f) AMMC projections of wildtype (grey) and VT059420-LexAGADfl>LexAop-RPR (purple) flies with antenna glued (N=9, average ΔF/F ± SEM, X-axis: seconds relative to flight induction). **(J)** Before and after plots of average ΔF/F during sound response in male and female flies (indicated above) with NP1046-Gal4 driving expression of GCaMP7f and exposure to initial saline (black), Dh44 pico injection (red), 170 SPI (light grey), 580 SPI (dark grey)(N=10, significance from two way repeated measures ANOVA with post-hoc Tukey’s test for individual comparisons: NS=P>0.05, *=P<0.05). **(K)** Before and after plots of average ΔF/F during sound response in male flies with NP1046-Gal4 driving expression of GCaMP7f and crispr mediated knock-out of Dh44R1 with exposure to initial saline (black), Dh44 pico injection (red), 170 SPI (light grey), 580 SPI (dark grey)(N=10, significance from repeated measures ANOVA with post-hoc Tukey’s test for individual comparisons: NS=P>0.05). **(L)** Quantification of HCR puncta labeling Dh44R1 expression in JO-B neurons in single optical sections compared between males and females and males expressing crispr mediated Dh44R1 knock out (indicated on x-axis) (N=8,significance from one-way ANOVA with post-hoc Tukey’s test for individual comparisons: ****=P<0.0001, NS=P>0.05).

Since the JO-As have the highest expression of HisCl1 (Figure 3D) and receive the most input from the MtAHNs (Cheong et al., 2024; Dorkenwald et al., 2022; Schlegel et al., 2024), we next investigated the impact of histamine signaling on sound evoked responses in JO-As. JO-As were stimulated with a 200Hz sine sound stimulus at a particle velocity of 1 mm/s, which is within the reported dynamic range of *Drosophila* hearing (Effertz et al., 2011), and their activity was recorded using GCaMP7f transients within the AMMC (Figure S3.2A). Pico-injection of histamine (Figure 3A) significantly increased JO-A sound-evoked responses which began to return to initial levels around 580 seconds post injection (SPI) (Figure 3B, Figure S3.2D). Pico-injection of saline alone did not cause any significant change in response (Figure S3.2B-C) and selectively knocking down HisCl1 receptor in the JO-As was sufficient to abolish the enhancement of sound evoked responses by histamine (Figure 3C, Figure S3.1A-C, Figure S3.2E). HisCl1 is a histamine-gated chloride channel which, under normal conditions, causes inhibition when active (Pantazis et al., 2008; Zheng et al., 2002). However, primary afferents often have higher internal chloride concentrations, thus opening a chloride channel can cause a form of inhibition called primary afferent depolarization (PAD) (Baden & Hedwig, 2010; Cattaert et al., 1994; Deschenes et al., 1976; Poulet & Hedwig, 2006; Wall, 1958). PADs are traditionally considered inhibitory and in line with this, we observed that longer bath application of histamine suppressed sound evoked responses in JO-As (Figure 3E, S3.3A-D). Chloride channel activation by PADs can also cause increases in intracellular calcium levels without mechanosensory input (Baden & Hedwig, 2010) and consistent with this, JO-As exhibit increases in initial intracellular calcium levels when HisCl1 is activated by histamine just prior to sound presentation (Figure 3D-Di). Overall, these results are consistent with a model in which histamine evokes PADs in JO-A auditory afferents.

We next sought to determine if the MtAHNs provide suppressive input to JO-As during flight. Since the natural context for the activation of the MtAHNs is flight (Cheong et al., 2024) and flight itself causes reafference (Mamiya & Dickinson, 2015), we imaged the activity of the JO-As during flight, but glued the antennae to prevent reafferent activation of the JO-As (Figure 3F). We then drove the expression of rpr to ablate the MtAHNs (validated by the loss of histamine labeling signal in the AMMC; Figure 3G, S1.4B) to determine if the MtAHNs provide a suppressive signal to the JO-As during flight. When the antennae are glued and the MtAHNs are intact, the induction of flight causes little to no change in activity of JO-As (Figure 3H-I). However, The JO-As of flies without MtAHNs showed a robust increase in activity following the induction of flight (Figure 3H-I). This implies both that during flight JO-As are inhibited by the MtAHNs and receive excitatory input from other circuits such as ascending leg proprioceptors that extend from the ventral nerve cord to synapse upon JONs and other AMMC cell types(S.-Y. J. Lee et al., 2025).

Although the JO-Bs receive less synaptic input from the MtAHNs relative to the JO-As, they express Dh44R1 at a higher level relative to the other JON subtypes (Figure 2E). To determine the effects of Dh44 on the JO-Bs, we recorded the responses of JO-B neurons to 100Hz sound stimulation (Figure S3.4A) and used pico injection to deliver Dh44 (Figure 3A). Surprisingly, Dh44 has a sex-specific effect, causing a robust decrease in the sound-evoked responses of JO-Bs in males which did not dissipate after 10 minutes (Figure 3J, Figure S3.4B-D) and having no effect on the responses of JO-B neurons in females. To confirm that Dh44 directly suppresses JO-Bs in males, we selectively knocked down Dh44R1 in the JO-Bs which eliminates the effects of Dh44 pico-injection JO-B sound evoked responses (Figure 3K, S3.5). Separately analyzing HCR labeling of Dh44R1 mRNA in JO-Bs in males and females revealed that only male JO-Bs express Dh44R1 (Figure 3L). Thus, histamine and Dh44 respectively decrease JO-A and JO-B sound responses, however the effects of each neurotransmitter occur over separate timescales and Dh44 has a male specific effect.

### AMMC B1 neurons downstream of the MtAHNs are inhibited via Dh44R1

We next explored the effects of the MtAHNs on second order auditory neurons in the AMMC. In the FlyWire EM dataset (Dorkenwald et al., 2022; Schlegel et al., 2024), a subset of AMMC B1s are the top downstream targets of the MtAHNs in the AMMC (Figure 1B, Figure S1.3F). Generally, the AMMC B1 neurons are responsible for conspecific song feature detection (Baker et al., 2022; Kamikouchi et al., 2009; Lai et al., 2012; Tootoonian et al., 2012; Vaughan et al., 2014). However, the AMMC B1 neurons are a diverse cell type and their downstream targets are not limited to courtship neural circuits (Dorkenwald et al., 2022; Schlegel et al., 2024), thus they may play a more generalized role in auditory stimulus detection (A. W. Azevedo & Wilson, 2017). The AMMC B1s do not appear in the reporter lines for HisCl1 or Ort (Figure 1C-D); thus, we reasoned they might express one of the Dh44 receptors. The characteristic contralateral branching pattern of the AMMC B1s was not evident in a Dh44R2 reporter line (Figure 4A), however this characteristic morphological trait was present in a Dh44R1 line (Figure 4B) and became more evident with stochastic labeling of this line (Figure 4Bi) using the multi-color flip out technique (MCFO;(Nern et al., 2015)). Next, we used FlyLight (Jenett et al., 2012) to identify R47F09-Gal4 as a driver line expressed primarily by AMMC B1 neurons and verified that the AMMC B1 neurons in this line express *Dh44R1*, but not *HisCl1* or *Ort*, using HCR (Figure 4D-Dii, F-Fii). We furthermore used MCFO to stochastically label the AMMC B1 neurons in R47F09-Gal4 and aligned the LM images in NeuronBridge (Clements et al., 2024) to identify matches in the FlyWire EM dataset. The top EM matches for each of the single AMMC B1 clones from R47F09-Gal4 were amongst the top targets of the MtAHNs in the AMMC based on the synapse predictions from FlyWire (Figure S1.1F, S4.1). The MtAHNs have relatively low synapse counts onto individual partners in the AMMC, which may reflect paracrine signaling by Dh44. Furthermore, some of the AMMC-B1 neurons identified in FlyWire (such as CB3719, Figure S4.1i) did not have strong matches to clones from our MCFO experiments indicating that R47F09-Gal4 does not include all AMMC-B1 neurons downstream of the MtAHNs. Additionally, there were additional soma with *Dh44R1* HCR labeling near those of the R47F09 AMMC-B1s. Thus, the R47F09-Gal4 includes some but not all *Dh44R1*+ AMMC B1 neurons. Some AMMC B1 neurons express *fruitless* indicating they may be sexually dimorphic (Cachero et al., 2010; Zhou et al., 2015b), however visual comparison of the R47F09-Gal4 AMMC B1 morphology revealed no obvious differences between males and females (Figure 4C), and there were no differences in Dh44R1 and R2 labeling intensity for these neurons between males and females (Figure 4E-Ei). Thus, AMMC B1 neurons are downstream of the MtAHNs and many AMMC B1 neurons express the Dh44R1 in both males and females.

**Figure 4.**
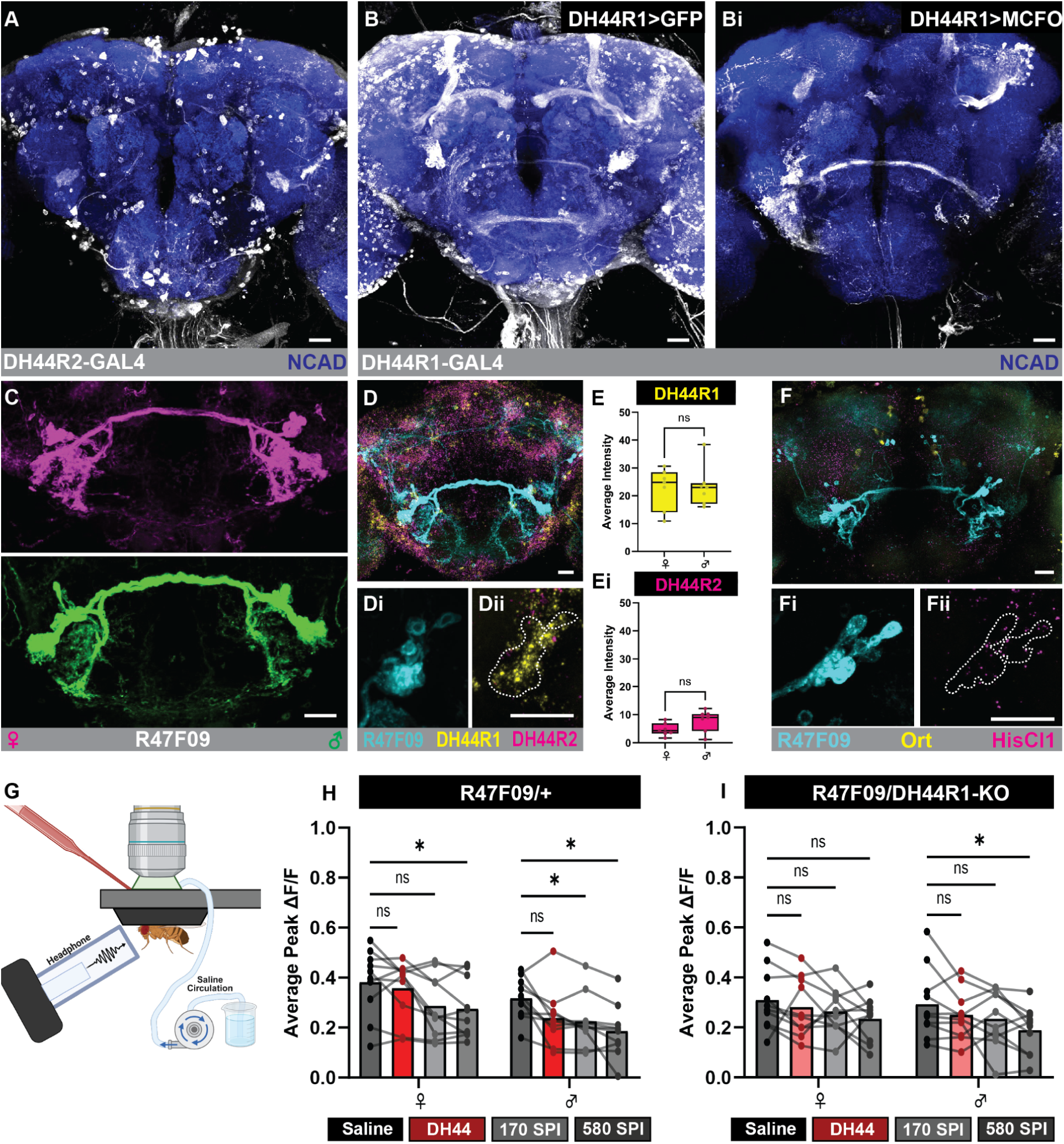
Dh44 suppresses sound evoked responses in AMMC B1 neurons. **(A)** GFP expression driven by the Mi{Trojan-Gal4.2}Dh44-R2[MI09411-TG4.2] line (white) with NCAD as a neuropil marker (blue). **(B)** GFP expression (white) and **(Bi)** MCFO mediated stochastic labeling (white) driven by the TI{GFP[3xP3.cLa]=CRIMIC.TG4.2}Dh44-R1[CR00985-TG4.2] line (white) with NCAD as a neuropil marker (blue). **(C)** GFP expression driven by R47F09-Gal4 in male (green) and female (magenta) AMMC B1 neurons. **(D)** Full brain GFP expression driven by R47F09-Gal4 (cyan) with HCR labeling for Dh44R1 (yellow) and Dh44R2 (magenta). **(Di)** Cell body image stack of AMMC B1 neurons in the R47F09-Gal4 line (cyan) with **(Dii)** their location in reference to the Dh44R1 (yellow) and Dh44R2 (magenta) labeling indicated via white dotted lines. **(E-Ei)** Boxplot of HCR labeling fluorescence in males and females for **(E)** Dh44R1 and **(Ei)** R2 in cell bodies of AMMC B1s in R47F09-Gal4 (N=7, significance from unpaired t-tests: NS=P>0.05, error bars=STDEV). **(F)** Full brain GFP expression driven by R47F09-Gal4 (cyan) with HCR labeling for Ort (yellow) and HisCl1 (magenta). **(Fi-Fii)** Cell body image stack of the AMMC B1s in the R47F09-Gal4 (cyan) with HCR labeling for Ort (yellow) and HisCl1 (magenta), cell body location indicated with white dotted lines. **(G)** Schematic of experimental setup for recording sound responses in AMMC B1s with Dh44 pico-injection, created with BioRender.com. **(H-I)** Before and after plots of female and male average ΔF/F during sound presentation in initial saline recording (black), 40μM Dh44 pico injection (red), 170 SPI (light grey), and 580 SPI (dark grey) in **(H)** R47F09>GCaMP7f flies and **(I)** R47F09/Dh44R1-KO>GCaMP7f flies (N=10, significance from two-way repeated measures ANOVA with post-hoc Tukey’s test for individual comparisons: NS=P>0.05, *=P<0.05). Scale bars = 20μm.

To determine the impact of Dh44 on the sound evoked responses of AMMC B1 interneurons, we recorded the responses of R47F09-Gal4 AMMC B1 neurons to 100Hz frequency sound before and after Dh44 pico-injection (Figure 4G). Pico-injection of Dh44 caused a significant decrease in the responses of R47F09-Gal4 AMMC B1 neurons that was sustained for at least 10 minutes (Figure 4H). The suppressive effects of Dh44 on AMMC B1 interneurons was present in males and females, however the decrease was stronger in males (Figure 4H, Figure S4.2B). Bath application of Dh44 also induced a decrease in AMMC B1 sound evoked responses that was sustained in males, but not in females (Figure S4.2 E-Ei). This sex-specific difference may arise from the combined suppressive effects of Dh44 directly on AMMC B1 neurons and their presynaptic partners, the JO-Bs (Figure 3J) in males. Consistent with this hypothesis, driving a CRISPR-mediated knockout of Dh44R1 expression in the R47F09-Gal4 AMMC B1 neurons eliminated the suppressive effects of Dh44 on sound evoked responses in females, but only reduced and delayed the suppressive effects of Dh44 in males (Figure 4I, S4.2C, S4.3). Therefore, the Dh44 reduces sound-evoked responses of AMMC B1 neurons in both males and females, but has a more robust effect in males where it exerts both pre-and postsynaptic modulation.

### Histamine and Dh44 release by the MtAHNs affect copulation success

Given the role of the AMMC-B1s in processing conspecific courtship song (Baker et al., 2022; Kamikouchi et al., 2009; Lai et al., 2012; Tootoonian et al., 2012; Vaughan et al., 2014), we investigated the impact of either histamine or Dh44 transmission by the MtAHNs on courtship behavior. We disrupted either histamine synthesis or neuropeptide transmission in the MtAHNs of male flies using the MtAHN splitGal4 to express HDC-RNAi (Figure S5.1D) or StacL-RNAi (Figure S5.1C-Cii) respectively. Male courtship behavior was scored based on completion of each step within the courtship sequence of *Drosophila melanogaster* (orientation, tapping, wing extension, licking, mounting, copulation (Spieth, 1974)) within an arena containing a single female (Figure 5A-B). Although knocking down either HDC or StacL in the MtAHNs does affect the likelihood that a male will attempt to court (Figure S5.1A-B), it does result in a significant decrease in successful copulation (Figure 5C). Furthermore, there was an increase in courtship attempt failures at the wing extension stage (Figure 5D) which is the phase in which courtship song is completed by the male (Billeter et al., 2006). Thus, the MtAHNs likely play a role in auditory processing during courtship behavior.

**Figure 5.**
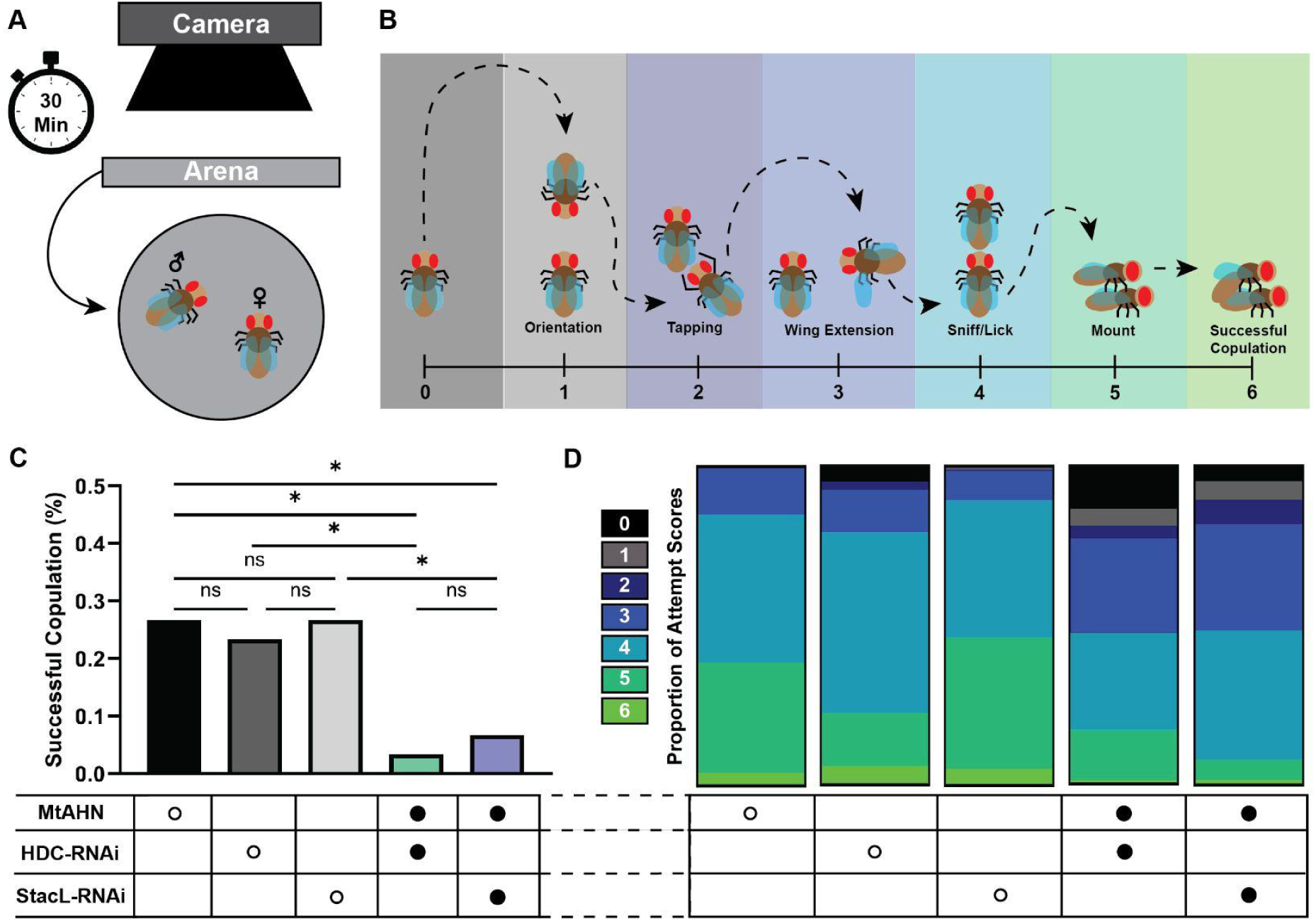
Both histamine and Dh44 release from the MtAHNs are necessary for successful copulation. **(A)** Schematic of experimental set up and **(B)** scoring parameters for courtship stage completion. **(C)** Bar graph of percentage of flies from each genotype who completed successful copulation with genotype indicated below (N=30, significance from one-way ANOVA with post-hoc Tukey’s test for individual comparisons: NS=P>0.05, *=P<0.05). **(D)** Proportions of scores from total attempts of each genotype, score value indicated with color key and genotype indicated in table below.

### Dh44 Co-transmission correlates with courtship modality across species

The AHNs are conserved among many arthropods, but their morphology varies, in some cases reflecting species-specific body plan and sensory ecology (Chapman et al., 2017; Maurer et al., 2019; Rieger & Harzsch, 2008; Skiebe et al., 1990). We surmised that co-transmission of Dh44 (or other neuropeptides) by the MtAHNs may also vary based on species specific behavioral ecology. Since the JO-Bs and AMMC B1s are associated with processing of courtship song, we hypothesized that Dh44 co-transmission by the MtAHNs is correlated with the production of auditory signals during courtship. The AHNs survive metamorphosis in other insect species (Bradley et al., 2016), yet larvae do not produce courtship displays, so we hypothesized that larval MtAHNs would not co-transmit Dh44. We confirmed that the MtAHNs are present in the larval CNS of *Drosophila melanogaster* with histamine antibody labeling, but that they do not express either *Dh44* (Figure 6A) or *StacL* (Figure S6.2Ai). Thus, the larval MtAHNs are likely not peptidergic, consistent with the hypothesis that Dh44 transmission is correlated with auditory courtship behaviors.

**Figure 6.**
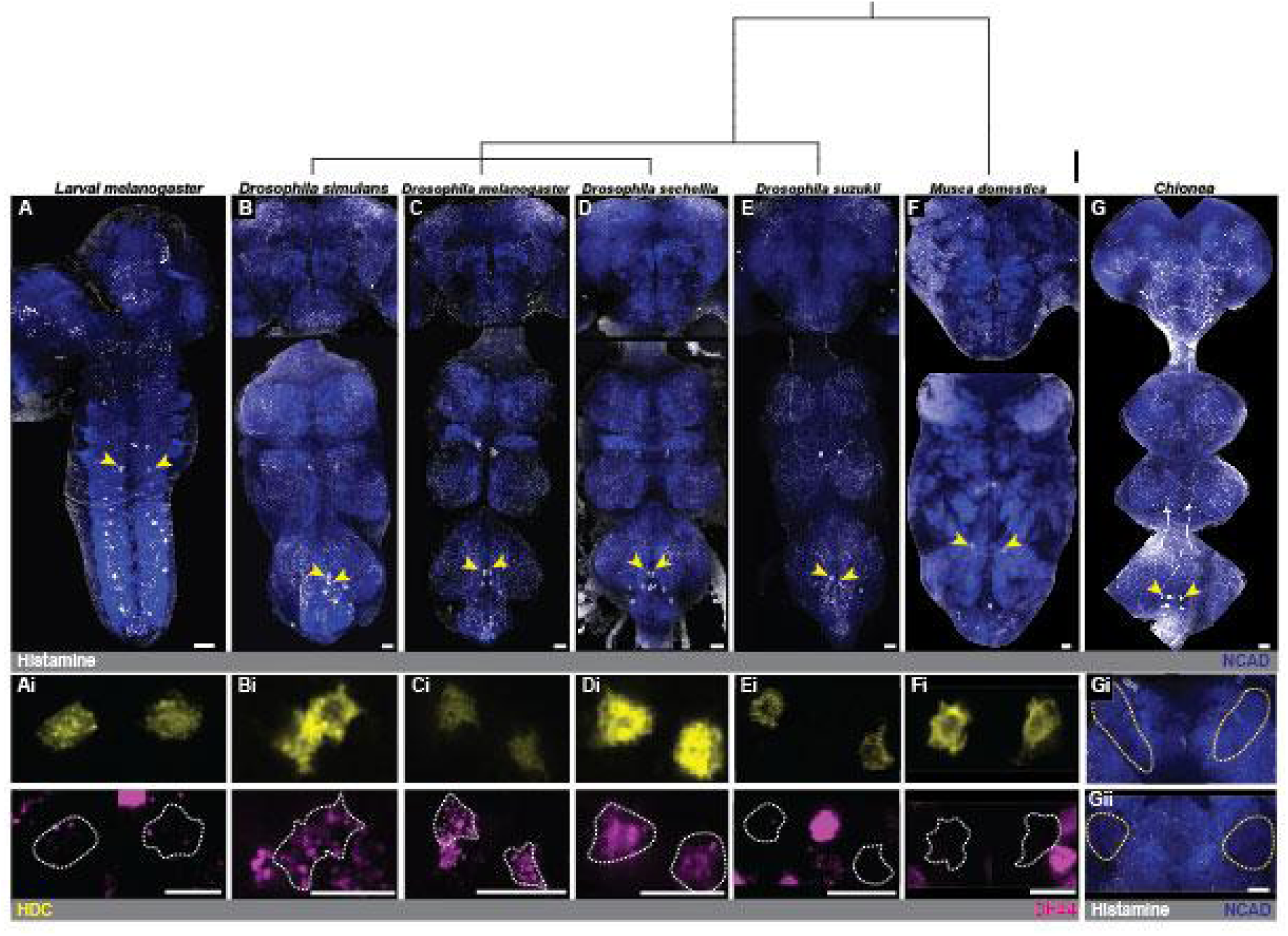
The MtAHNs survive metamorphosis and are conserved in other species of flies but only co-transmit Dh44 in singing drosophilids. (A-G) Histamine antibody labeling (white) with NCAD as a neuropil marker (blue) in the full CNS of several *Drosophila* sp., *M. domestica,* and snow flies from the genus *Chionea*. MtAHN cell bodies indicated with yellow arrows and species indicated above with phylogenetic tree showing evolutionary relatedness of each species (scale bar = ∼10 million years). **(Ai-Fi)** MtAHN cell bodies with HCR labeling for HDC (yellow) and Dh44 (magenta) in species indicated above. MtAHN cell bodies indicated with white dotted lines. **(Gi-Gii)** Histamine antibody labeling (white) in AMMC (yellow dotted line) of *Chionea* with NCAD as a neuropil marker (blue). Scale bars = 10μm.

We next hypothesized that the MtAHNs would only co-transmit Dh44 and histamine in *Drosophila* species that use their wings to generate courtship song. *D. simulans* and *D. sechellia* are closely related to *D. melanogaster* and their courtship displays include the generation of songs produced by wing vibration (Cobb et al., 1989; Ewing & Bennet-Clark, 1968). After confirming the presence of MtAHNs using antibody labeling of histamine in *D. simulans* (Figure 6B) and *D. sechellia* (Figure 6D), we used HCR to label for *HDC* and *Dh44* in the full central nervous systems of both species (Figure S6.1B,C). Consistent with our hypothesis, HCR labeling for *HDC* and *Dh44* in the MtAHNs overlapped in both *D. simulans* (Figure 6Bi) and *D. sechellia* (Figure 6Di), as did the expression of *StacL* (Figure S6.2Ci,Di). Therefore, co-transmission of histamine and Dh44 by the MtAHNs was conserved across three species of *Drosophila* that use auditory courtship.

Next, to determine if the MtAHNs co-transmit histamine and Dh44 in species of flies that court using non-auditory cues, we turned to *D. suzukii*, a closely related species to *D. melanogaster* that uses a purely visual wing display during courtship (Revadi et al., 2015). After confirming the presence of the MtAHNs using antibody labeling of histamine (Figure 6E), we used HCR in the full central nervous system of *D. suzukii* (Figure S6.1D) and found no overlap of *HDC* and *Dh44* (Figure 6Ei) or *StacL* (Figure S6.2E) in the MtAHNs, indicating they are not likely peptidergic. Although this is consistent with Dh44 co-transmission being limited to auditory courtship behaviors, it is also possible that the loss of Dh44 in *D. suzukii* was a consequence of visually driven courtship behavior, and that Dh44 co-transmission represents the basal state for the MtAHNs. Therefore, we investigated whether the MtAHNs co-transmit histamine and Dh44 in a species of fly with no known courtship behavior, *Musca domestica* (ter Haar et al., 2023). After confirming the presence of the MtAHNs (Figure 6F) in *M. domestica* using antibody labeling, we used HCR to label for the mRNA transcripts of *HDC* and *Dh44* in the full central nervous system (Figure S6.1E) and found no overlap in the MtAHNs (Figure 6Fi). Thus, co-transmission of histamine and Dh44 by the MtAHNs is correlated with the use of auditory signals for courtship.

Histaminergic processes of the MtAHNs in the AMMC are conserved among the aforementioned species consistent with a role in modulating auditory responses during flight (Figure 6). However, innervation of sensory neuropil by the ascending histamine neurons is known to vary in the olfactory system of the Lepidoptera based on sensory ecology (Chapman et al., 2017). We therefore sought to determine if the MtAHNs in the wingless snow fly (Chionea spp.) innervate the AMMC, as the lack of wings in this species changes the reafferent stimulation that the Johsnton’s organ receives during locomotion. To this end, we labeled the central nervous system of *Chionea* for histamine and determined that while the MtAHNs in this species ascend to the brain, but do not seem to enter the AMMC (Figure 6G). Therefore, MtAHN morphology and co-transmission vary with the sensory ecology and behavioral repertoire across species.

## Discussion

Reafferent stimulation can impact processing of sensory information within separate behavioral contexts(Jékely et al., 2021; Von Holst & Mittelstaedt, 1950). Predictive motor circuits must therefore provide differential signaling to distinct sensorimotor circuits within timescales specific to each behavioral outcome. *Drosophila melanogaster* beat their wings at a frequency of∼200Hz (Lehmann & Dickinson, 1997) which causes mechanosensory reafference in the JO-As and JO-Bs (Mamiya & Dickinson, 2015). Although flight and courtship circuits receive similar mechanosensory input their circuit architecture is very different. The giant fiber escape circuit is very direct, using just one type of descending neuron to drive motor output through electrical synapses (Ache et al., 2019; Jezzini et al., 2018; Pézier et al., 2014; Von Reyn et al., 2017; Wyman et al., 1984). Conversely, the courtship song recognition circuit is more indirect, with multiple levels of sensory integration preceding motor output commands (Baker et al., 2022; Tootoonian et al., 2012; Zhou et al., 2015b). This circuit architecture mirrors the nature of these behaviors, as escape is fast to avoid predation (Von Reyn et al., 2017) and courtship is slow and involves many stages to accomplish successful copulation (Billeter et al., 2006). Thus, it stands to reason that the regulation of these circuits would occur via signaling mechanisms that differ in their time course.

Using the MtAHNs as a model predictive motor circuit, we demonstrate that co-transmission of fast and slow neurotransmitters can enable a single network component to differentially modulate distinct sensory sub-circuits. The MtAHNs become active immediately before the onset of flight (Cheong et al., 2024) and project to the AMMC (Figure 1A) where they synapse upon a collection of auditory neurons including primary auditory afferents and secondary auditory interneurons. We show that in *D. melanogaster*, the MtAHNs co-transmit histamine and Dh44, and that different auditory neuron types express receptors for either transmitter. The expression of the HisCl1 receptor by the JO-A sensory afferents (Figure 3D) supports immediate and short lived PAD-like responses and ablating the MtAHNs revealed latent excitatory synaptic input to the JO-As (Figure 3), consistent with PADs providing inhibition during the articulation of behavior. The JO-Bs and the AMMC-B1s express the Dh44R1 receptor (Figure 3E, 4D-E) and Dh44 causes a delayed, yet sustained reduction in sound-evoked responses in both neuron types (Figure 3J, 4H, S3.4D, S4.2B,E). Unfortunately, our attempts to record the MtAHN activity from the VNC during courtship required physiological preparation that proved too disruptive to enable optogenetic triggering of wing movement by courtship neurons, so we cannot make any claims as to the nature of MtAHN activity during courtship. However, loss of either histamine or Dh44 by the MtAHNs reduced copulation success (Figure 5) and co-transmission by the MtAHNs is only present in species that produce courtship song (Figure 6), consistent with a model in which each transmitter impacts auditory processing in distinct behavior contexts. Thus, the MtAHNs send predictive signals to the AMMC during flight (Cheong et al., 2024), yet differences in receptor expression between auditory neurons could provide a means to differentially impact the behavioral circuits within which they are integrated.

As locomotor strategies change over the course of evolution to match biophysical constraints and sensory ecology of a given species, corollary discharge circuits must also shift to provide appropriate predictions. The AHNs are conserved across many arthropods, however their morphology changes depending on body plan and species specific behavior (Chapman et al., 2017; Maurer et al., 2019). In moths and caddisflies, the AHNs have additional processes that project into the antennal lobe (Chapman et al., 2017). In the tobacco hawkmoth *Manduca sexta* the AHNs act within a predictive motor circuit modulating the olfactory system to aid in odor pulse tracking created by wing beating during flight (Bradley et al., 2016; Chapman et al., 2018). The additional antennal lobe projections are not present in *Drosophila* or butterflies, suggesting that species-specific differences in flight mechanics or sensory ecology alter the consequences of reafference from wing-beating (Chapman et al., 2017). While the AHNs project into the AMMC in Lepidoptera(Chapman et al., 2017), Drosophilids (Figure 6B-E), and *M. domestica* (Figure 6F), they are lost in the wingless snow fly (Figure 6G), demonstrating that the functional properties of predictive motor circuits vary with body plan and locomotion strategy. Furthermore, species specific variation in co-transmission in correlation with use of courtship song (Figure 6) suggests that there are likely many parameters of CDIs in addition to morphology that vary over the course of evolution.

The AMMC receives input from mechanosensory neurons in the Johnston’s organ (JO) that respond to auditory, wind, and gravity signals (Boyan et al., 2023; Burkhardt & Gewecke, 1965; Gewecke, 1970; Ishikawa et al., 2020; Kladt & Reiser, 2023; Mukunda & Sane, 2025; Patella & Wilson, 2018; Roy Khurana & Sane, 2016; Sun et al., 2009). In some insects, the AMMC processes both flight and conspecific communication signals (Baker et al., 2022; Dreller & Kirchner, 1993; Loh et al., 2023; Mamiya & Dickinson, 2015; Matsuo et al., 2014; Pennetier et al., 2010; Roy Khurana & Sane, 2016; Suver et al., 2023; Tootoonian et al., 2012; Zhou et al., 2015b), whereas in other species like hawk moths, locusts and blowflies the AMMC’s sole purpose seems to be maintaining flight stability(Budick et al., 2007; Burkhardt, 1960; Burkhardt & Gewecke, 1965; Gewecke, 1970; Loh et al., 2023; Mamiya & Dickinson, 2015; Mukunda & Sane, 2025; Roy Khurana & Sane, 2016). We found that the AHNs project into the AMMC among all flying species tested; *D. simulans*, *D. sechellia*, *D. suzukii*, and *M. domestica*, but not in the wingless *Chionea* sp.(Figure 6B-G). This suggests that the AHNs may serve a conserved role modulating the mechanosensory input to the JO during flight. In *Drosophila melanogaster* histaminergic input to the AMMC is in part directed at the subtype of JONs that give the most direct input to the giant fiber neurons(Figure 2D) (Baker et al., 2022; Jezzini et al., 2018; Pézier et al., 2014). Many arthropods use giant fiber neurons to drive escape behavior related to sensory stimuli derived from approaching predators (Belosky & Delcomyn, 1977; Farley & Milburn, 1969; Herberholz, 2022; Seabrook, 1971; Wyman et al., 1984). Giant fiber escape circuits also tend to have a CDC associated with them which prevents reafferent stimuli from desensitizing mechanosensory circuits (Crapse & Sommer, 2008; Delcomyn, 1977; Schöneich & Hedwig, 2015). In crickets and cockroaches, CDIs inhibit sensory afferent input to the giant fiber circuit during behaviors that produce reafference such as singing, walking, and flying (Belosky & Delcomyn, 1977; Delcomyn, 1977; Schöneich & Hedwig, 2015). In the flying locust, sensory input to the giant fiber neurons is reduced through primary afferent depolarization of wind sensitive afferents (Boyan, 1988). Thus, CDIs often inhibit mechanosensory afferents that synapse onto the giant fiber escape neurons, often using classic IPSPs or PADs. Although the mechanisms of synaptic inhibition differ across these model systems, they all serve to regulate the sensitivity of mechanosensory afferents to allow for appropriately timed escape behavior. Therefore, it is possible the MtAHNs in *Drosophila melanogaster* elicit PADs-like responses in JO-As to maintain the sensitivity of these neurons for appropriate escape response. However, in *Drosophila melanogaster* flight reafference impacts more than just the JO-As and the giant fiber escape circuit, which presents the need for differential signaling from the MtAHNs.

Predictive motor circuits are essential for animals to process reafference generated by their own movement (Crapse & Sommer, 2008; Delcomyn, 1977; Ford & Mathalon, 2004; Poulet & Hedwig, 2006; Schöneich & Hedwig, 2015). However most behaviors cause diverse reafferent stimulation which impacts multiple sensorimotor circuits. How is this complex reafference processed within the CNS to provide an accurate comparison of self vs. environment derived stimuli? Here we use the MtAHNs as a model for CDIs tasked with processing complex reafference. We found co-transmission of fast and slow neurotransmitters provides a mechanism for differential processing of auditory stimuli and that it is conserved among invertebrates with similar behavioral ecology. Thus, co-transmission is likely co-opted by neurons which process differential sensory stimuli to aid in formulating an accurate comparison of self vs. environmental stimuli.

## Materials and Methods

### Fly lines

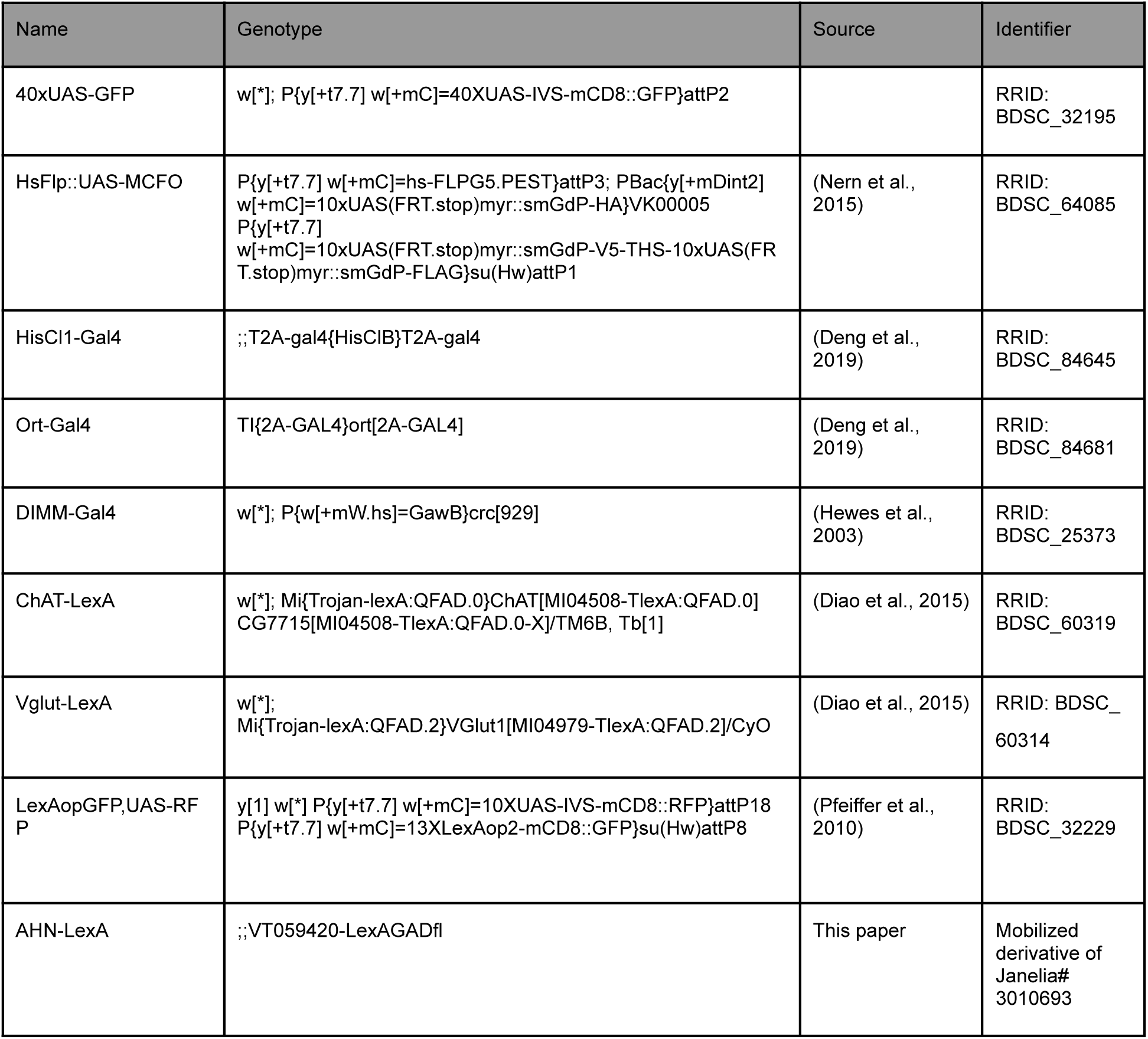
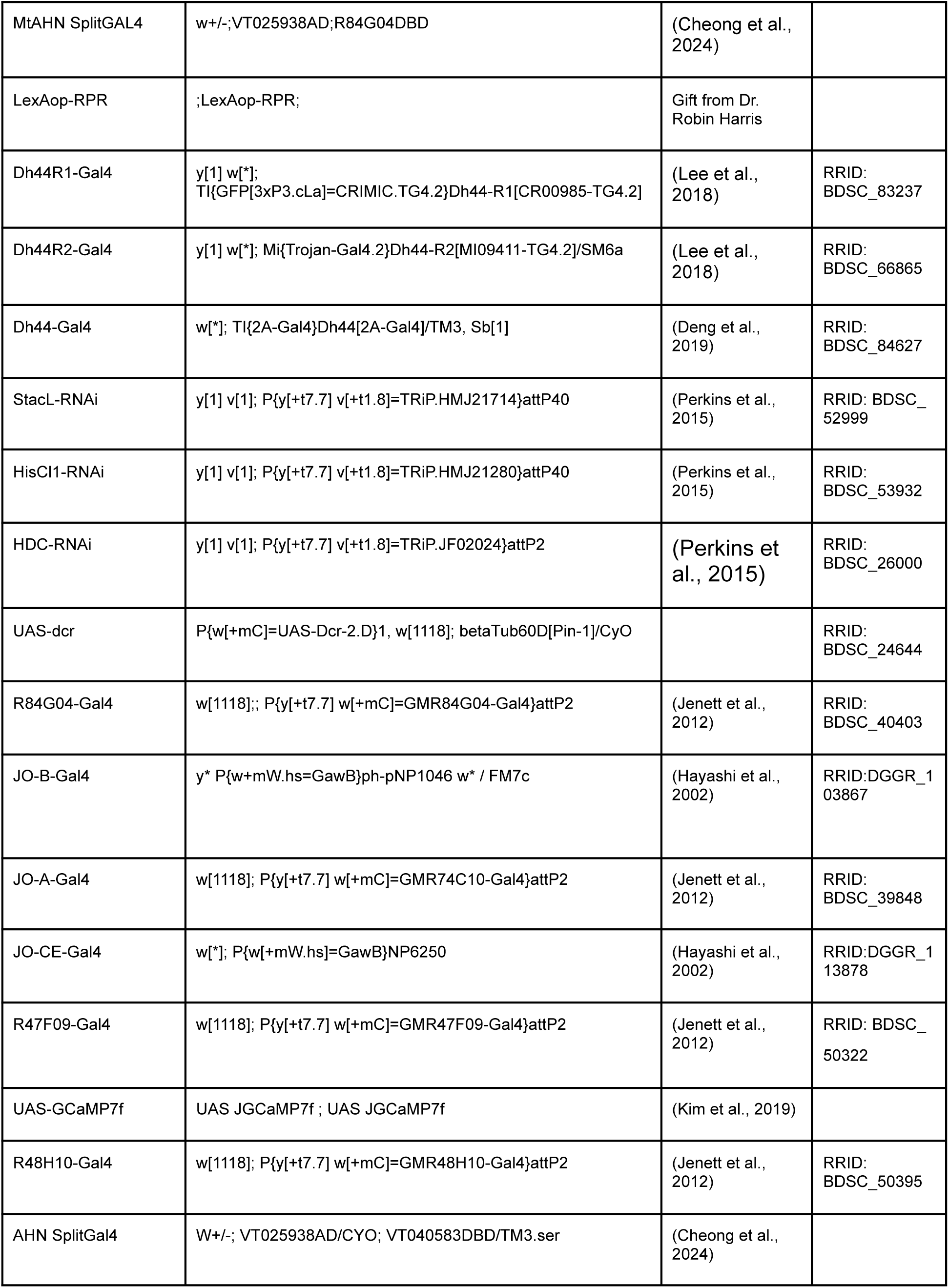
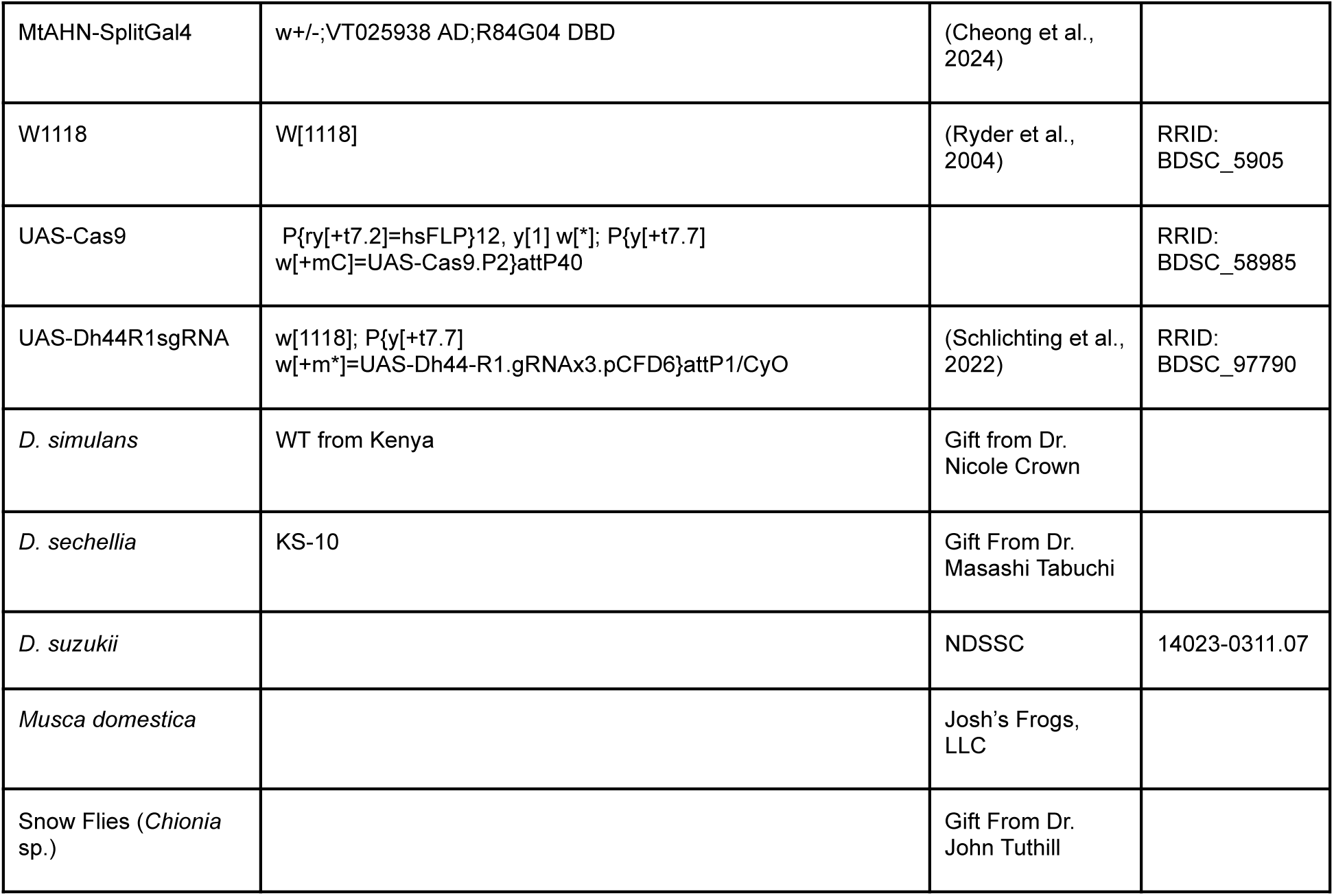

### Fly Husbandry

All fly stocks were raised on standard cornmeal/agar/yeast media at 24-25℃ on a 12:12 light dark cycle at 60% humidity. Flies used for Ca^2+^ imaging experiments were housed in mixed cultures and aged 4-9 days old before imaging.

### Connectomic Analysis

The MtAHNs were previously identified by (Cheong et al., 2024) in the FlyWire segmentation of the Full Adult Fly Brain (FAFB)(Dorkenwald et al., 2022; Schlegel et al., 2024) and the Male Adult Nerve Cord (MANC)(Marin et al., 2023; Takemura et al., 2024) and Female Adult Nerve Cord datasets(Azevedo et al., 2024; Phelps et al., 2021). The downstream connectivity of the MtAHNs in the AMMC in v783 of FAFB was determined by filtering the “connections_princeton_no_threshold” synapse table (Yu et al., 2025) for rows where a MtAHN was the “pre_root_id” and AMMC was the “neuropil”. The cell type (or hemibrain type if a cell type was not available) of the downstream partner was identified by matching the partner’s ID to a published annotation table (Schlegel et al., 2024). A list of connectivity for the MtAHNs top downstream targets in MANC is provided in Table S1 and for the AMMC in FAFB is provided in Table S2. AMMC-B1s were identified anatomically (Baker et al., 2022; Lai et al., 2012; Vaughan et al., 2014; Zhou et al., 2015) and from community annotations in FlyWire (Dorkenwald et al., 2022; Schlegel et al., 2024). Downstream connectivity of the MtAHNs in v1.2.1 of MANC was queried with the “fetch_adjacencies” function of the neuprint-python Python package (Berg & Schlegel, 2022). Cell types were previously published in MANC (Marin et al., 2023). Connectivity strength was measured by the number of synapses with the MtAHNs.

### Immunocytochemistry and Immunohistochemistry

Adult flies were anesthetized on ice and dissected in external saline, full central nervous systems were removed and fixed in 4% paraformaldehyde in PBS for 30 minutes to 2 hours depending on the primary antibody (for exact times and temperatures refer to table 2). Larvae were removed from food using 30% sucrose in diH2O and dissected in external saline and fixed in 4% paraformaldehyde for 30 minutes at RT. For histamine labeling in *Drosophila*, the samples were first pre-fixed in a 4% carbodiimide in PBS solution for 2 hours on ice (for *M. domestica* and snow flies this was extended to 4 hours), and then post-fixed in 4% paraformaldehyde in PBS for 30 minutes at RT (Chapman et al., 2017; Dacks et al., 2010). Samples were washed 4×15 minutes in PBS with 0.5% triton x-100 then blocked for 1 hour at RT with agitation. Primary antibodies were applied (see table 2 for concentration) in blocking solution with PBSAT with 5% Triton X-100 for 48 hours at 4℃ with agitation. Samples were washed and blocked as on day 1, secondary antibodies were added to blocking solution at concentrations according to table 2 and spun down at 4℃ in a microcentrifuge at 15,000 RPM. Secondary antibody solution was applied to the samples for 48 hours at 4℃ with agitation. Samples were then washed 2×15 minutes in PBST and 2×15 minutes in PBS at RT with agitation. Samples were cleared in an ascending glycerol series (40%, 60%, 80%) for 10 minutes each and then mounted on well slides using Vectashield (Vector laboratories, cat#H-1000-10) or Everbrite (Biotium, cat#23001).

**Table 2.** Primary and secondary antibody concentrations, blocking agents, and fixation time and temperature. “*” indicates that values can be varied based on blocking and fixation times needed for other antibodies being used. “#” indicates both prefixation and fixation steps are required, see above for details.

| Antigen | Host | Concentration | Blocking Agent | Fix Time (Minutes) and Temp | Source |
| --- | --- | --- | --- | --- | --- |
| AstA | Mouse | 1:250 | 3% NGS | 120 at RT | DSHB cat#5F10 |
| AstC | Rabbit | 1:1000 | 3% NGS | 240 at 4°C | (Veenstra et al., 2008) |
| CRZ | Rabbit | 1:1000 | 3% NGS | 60 at RT | (Veenstra & Davis, 1993) |
| Dh31 | Rabbit | 1:500 | 2% BSA | 120 at 4°C | (Park et al., 2008) |
| Dh44 | Rabbit | 1:1000 | 2% BSA | 60 at RT | (Cabrero et al., 2002) |
| FMRFamide | Rabbit | 1:1000 | 5% NGS | 240 at 4°C | (Marder et al., 1987) |
| GABA | Rabbit | 1:500 | 2% BSA | 30 at RT | Sigma Cat#A2052 |
| GFP | Chicken | 1:1000 | * | * | Invitrogen Cat#PA1-9533 |
| Histamine | Rabbit | 1:1000 | 4% NGS | # | Immunostar Cat#22939 |
| Leucokinin | Rabbit | 1:250 | 3% NGS | 240 at 4°C | (Zhang et al., 2017) |
| MIP | Rabbit | 1:4000 | 2% BSA | 60 at 4°C | Gift from Wegener |
| N-sytAlpha | Rabbit | 1:500 | 4% BSA | 30 at RT | (Adolfson et al., 2004) |
| N-sytBeta | Rabbit | 1:500 | 4% BSA | 30 at RT | (Adolfson et al., 2004) |
| PDF | Mouse | 1:200 | 3% NGS | 60 at 4°C | DSHB cat#PDF C7 |
| RFP | Rabbit | 1:500 | 4% BSA | 30 at 4°C | Rockland cat#600-401-379 |
| SNPF | Rabbit | 1:2000 | 3% NGS | 120 at RT | (Johard et al., 2008) |
| DSK | Rabbit | 1:1000 | 3% NGS | 120 at RT | Jena cat#ABD-037 |
| TKK | Rabbit | 1:500 | 2% BSA | 30 at 4°C | (Veenstra et al., 2008) |
| TH | Mouse | 1:100 | 5% NGS | 60 at RT | Immunostar Cat#22941 |
| 5-HT | Rabbit | 1:5000 | 2% BSA | 30 at RT | Immunostar Cat#20080 |
| NCAD | Rat | 1:250 | * | * | DSHB Cat#DN-Ex #8 |
| HA-Tag | Rabbit | 1:300 | 4% BSA | 30 at 4°C | Cell signaling Cat#3724 |
| Chicken | Donkey | 1:1000 | * | * | Jackson cat#703-545-155 |
| Rabbit | Donkey | 1:500 | * | * | Invitrogen cat#A10040 |
| Mouse | Goat | 1:1000 | * | * | Invitrogen cat#A11030 |
| Rat | Donkey | 1:1000 | * | * | Abcam cat#150155 |
| Rabbit | Goat | 1:1000 | * | * | LifeTech cat#A11008 |
| V5-Tag | Mouse | 1:500 | * | * | BioRad<br>cat#MCA1360D550GA |

For multi-color flip-out (MCFO) samples heat shock was performed 72 hours prior to dissections in a water bath set to 37℃. For Dh44 Receptor lines the heat shock was unnecessary as they are highly expressed Gal4 lines, instead these flies were reared at 18℃ to limit flipping out the fluorescent protein tags.

### Visualizing GFP Expression in Appendages

Adult *Drosophila melanogaster* were anesthetized on ice and dissected in external saline. For antenna and proboscis imaging, whole heads were removed and mounted on a well slide in the Vectashield (Vector laboratories, cat#H-1000-10). Legs, wings, and halteres were removed and mounted in Vectashield (Vector laboratories, cat#H-1000-10) on a well slide. These samples were not fixed prior to mounting, and were scanned immediately to prevent losing GFP signal from tissue degradation.

### Hybridization Chain Reaction (HCR)

#### Reagents

RNA probes were synthesized by Molecular Instruments to match the transcripts of the genes of interest provided by NCBI accession numbers and assigned amplifier binding sequences (table 3) and stored at-20℃. Amplifier hairpins (Molecular Instruments) with either B1-546nm, B2-647nm, B3-647nm fluorescent labels were stored in the dark at-20℃. Hybridization, Probe Wash, and Amplification buffers (Molecular instruments) were stored at-20℃,-20℃, and 4℃ respectively.

**Table 3.**
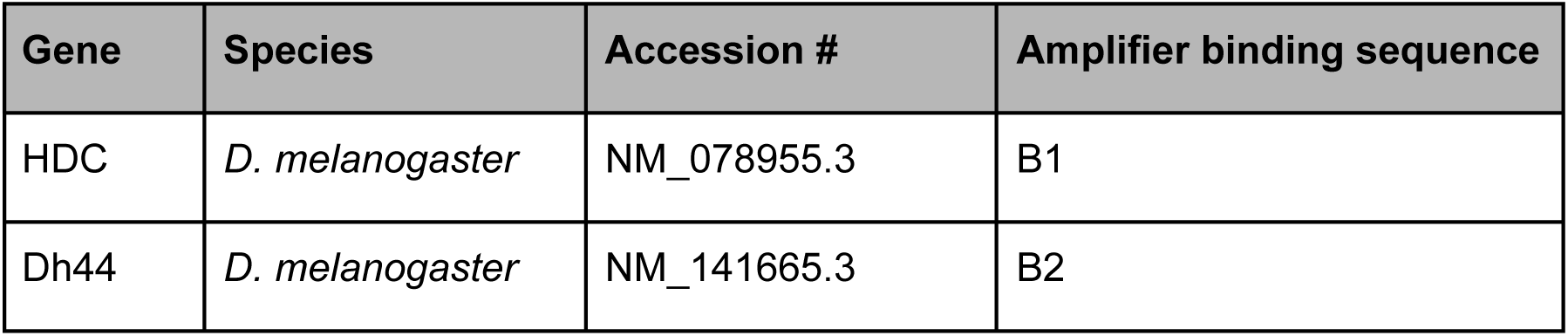
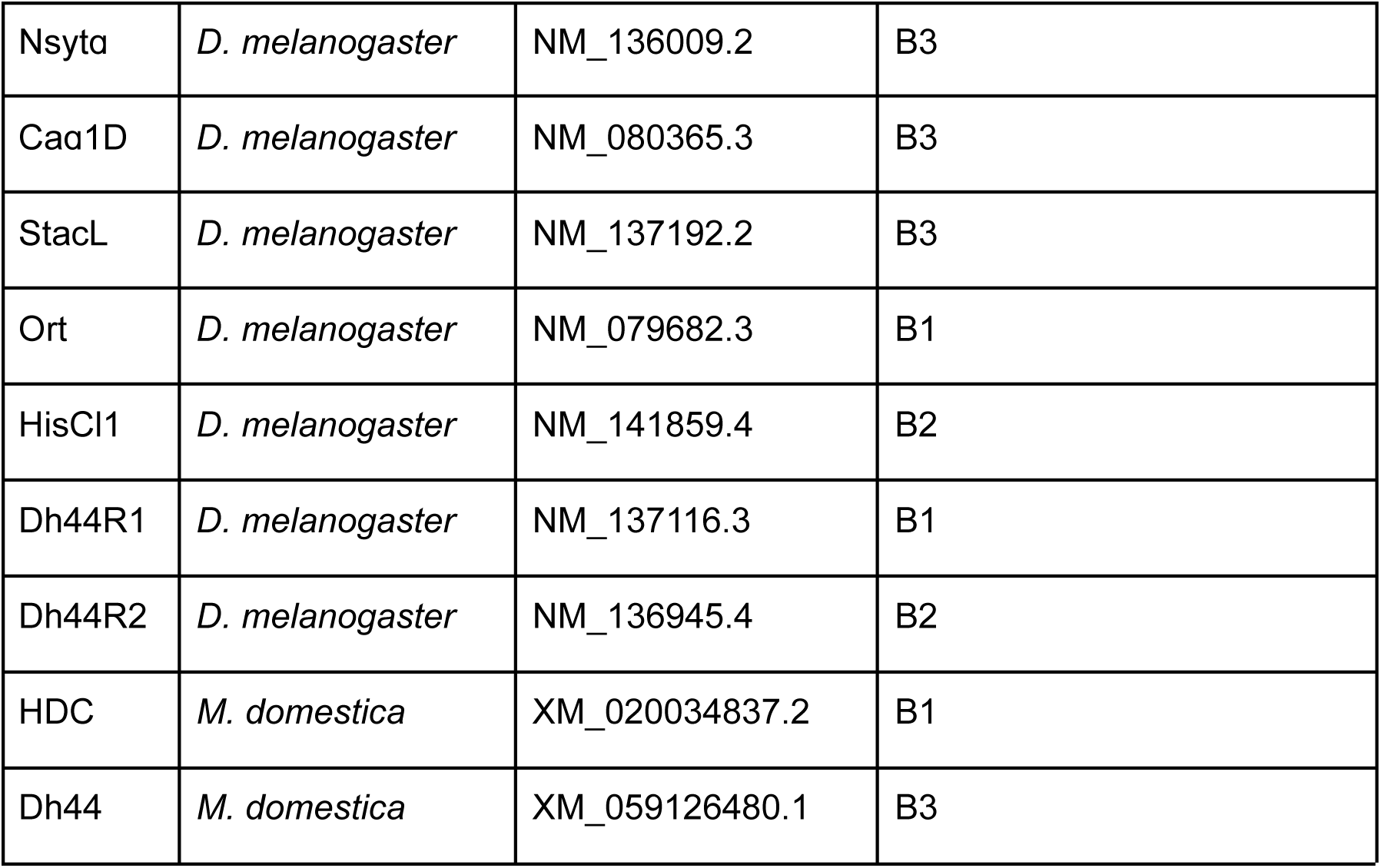
HCR probe generation information including gene, species, NCBI accession #s, and amplifier binding sequences.

#### Full CNS

Adult flies were anesthetized on ice and dissected in PBS. 3rd instar larvae were removed from food using 30% sucrose in diH2O and dissected in PBS. Samples were fixed at RT for 30 minutes in 4% paraformaldehyde in 0.01M PBS and then washed 4×15 minutes in PBST with 1% Triton x-100 at RT with agitation. Samples were pre-hybridized in hybridization buffer at 37℃ for 30 minutes with agitation. Probe solution was made by adding RNA Probes to hybridization buffer at a concentration of 1:250. Pre-hybridization solution was removed and samples were placed in probe solution at 37℃ overnight with agitation. Probe solution was then removed and samples were washed with probe wash buffer 4×15 minutes at 37℃ with agitation. Samples were then washed with 5xSSCT for 3×5 minutes at RT with agitation. Samples were pre-amplified in amplification buffer for 30 minutes at RT with agitation. Hairpins were snap cooled using a thermocycler by heating to 95℃ for 90 seconds and cooled to RT in a dark drawer for 30 minutes. Hairpin solution was prepared by adding 10ul of each hairpin to 500ul of amplification buffer. Pre-amplification solution was removed and replaced with hairpin solution in the dark at RT overnight with agitation. Hairpin solution was removed and samples were washed 2×5 minutes, 2×30 minutes, and 1×5 minutes in 5xSSCT in the dark at RT with agitation. Samples were cleared in an ascending glycerol series (40%, 60%, 80%) for 10 minutes each in the dark at RT with agitation. Samples were mounted in EverBrite (Biotium, cat#23001) or Prolong Glass (Invitrogen, cat#P36984). For all *Drosophila* species tested *D. melanogaster* probes were used and separate probes were generated for *M. domestica*.

#### Antennae

Adult *Drosophila melanogaster* were anesthetized on ice and skewered on a dissection pin through the wing muscle creating a “*Drosophila* kabob” which consisted of ∼5 flies per kabob. *Drosophila* kabobs were placed in 70% ethanol for 1 minute and lightly vortexed to remove oil. Kabobs were transferred to 4% paraformaldehyde to fix for 1 hour at RT. Post-fixation, kabobs were transferred to PBSAT with 5% Triton x-100 and washed 2×5 minutes at RT with agitation. Kabobs were removed and antennae were dissected and sectioned in PBS. Johnston’s organs (JO) were placed in PBSAT with 5% Triton as they were removed from the fixed flies. JOs were dehydrated in 100% methanol for 4×5 minutes and then rehydrated in a descending methanol in PBST with 1% triton series (75%, 50%, 25%) for 5 minutes each at RT with agitation. Samples were then washed 4×5 minutes with PBST with 1% Triton x-100 at RT with agitation. Hybridization, amplification, and mounting steps followed those of the full CNS prep except JO samples were kept in probe solution and hairpin solution for ∼48 hours each.

### Confocal imaging, Image processing, and Expression quantification

#### Scanning

Samples were scanned with an Olympus FV1000 BX61 (Shinjuku, Tokyo, Japan) confocal, using Fluoview FV1000 software with a 20xUPlanSApo, 40xUPlanFL-N or 60x PlanApo-N oil immersion objective. Appendages from the HisCl1-Gal4 samples and antennal HCR samples were scanned with an additional channel set to pick up cuticular autofluorescence for later subtraction.

#### Processing

Scans were processed using ImageJ to correct brightness and contrast. Appendage images were processed using the ImageJ plug-in “image calculator” to subtract the autofluorescence of the cuticle (Guan et al., 2018). For antennal HCR samples, scans were analyzed for expression levels using the “cell counter” plugin for imageJ to quantify HCR puncta in cells expressing GFP.

#### Quantification

For AMMC histamine immunofluorescence quantification an ROI was drawn according to the dimensions of the AMMC labeled with NCAD. The AMMC ROI was then used to measure the average intensity of the histamine IF in the image stack avoiding the labeling present in the SEZ. Another ROI drawn in an area of the image without histamine IF was used to measure the average intensity of the background. Background was subtracted from the AMMC IF to give the average intensity of the histamine labeling present in the AMMC. Quantification of receptor expression in JON samples was performed using the imageJ cell counter plugin. For each JO, cells expressing GFP were counted and annotated based on presence of overlap with puncta indicating expression of HisCl1, Dh44R1, or Dh44R2. Proportion of cells positive for expression of the gene of interest was calculated by dividing the total cells overlapping with puncta by the total cells expressing GFP. For quantification of Dh44R1 and Dh44R2 expression by the AMMC B1 neurons in the R47F09-Gal4 line, ROIs were drawn around the GFP signal in the cell bodies in each slice of the image. The cell body ROIs were then used to measure average intensity of HCR labeling for Dh44R1 and Dh44R2. An additional ROI was drawn in an area of the image without HCR labeling and used to measure the background intensity in the channel of interest. The average intensity of the background for each slice was subtracted from the intensity of the HCR labeling. The products of the background subtraction from each slice were averaged together and then the averages from the left and right cell bodies of each brain were averaged together.

### RNA Sequencing

#### Expression checks

Neurons of interest were isolated by expressing the dual fluorescent reporter 10XUAS-IVS-nls-tdTomato; 10XUAS-unc84-2XGFP with the driver R84G04-Gal4 and then manually picking the fluorescent neurons from dissociated VNC tissue. Prior to the sorting process, each driver/reporter combination was ‘expression checked’ to determine if the marked cells were sufficiently bright to be sorted effectively and if there was any off-target expression in neurons other than those of interest. Drivers that met both these requirements were used in sorting experiments as described below.

#### Sorting of fluorescent-labeled neurons

*Drosophila* adults were collected daily as they eclosed, and aged 3-5 days prior to dissection. For each sample, 60-100 VNCs were dissected in freshly prepared, ice cold Adult Hemolymph Solution (AHS, 108 mM NaCl, 5 mM KCl, 2 mM CaCl_2_, 8.2 mM MgCl_2_, 4 mM NaHCO_3_, 1 mM NaH_2_PO_4_, 5 mM HEPES, 6 mM Trehalose, 10 mM Sucrose). The VNCs were transferred to a 1.5 ml Eppendorf tube containing 500 microliters 1 mg/mL Liberase DH (Roche, prepared according to the manufacturer’s recommendation) in AHS, and digested for 1 hr at room temperature. The Liberase solution was removed, and the tissue washed three times with ice cold AHS. The final wash was removed completely and 400 microliters of AHS+2% Fetal Bovine Serum (FBS, Sigma) were added. The samples were gently triturated with a series of fire-polished, FBS-coated Pasteur pipettes of descending pore sizes until the tissue was homogenized, after which the tube was allowed to stand for 2-3 min so that the larger debris could settle.

For hand sorting, the cell suspension was transferred to a Sylgard-lined Glass Bottom Dish (Willco Wells), leaving the debris at the bottom of the Eppendorf tube, and distributed evenly in a rectangular area in the center of the plate with the pipet tip. The cells were allowed to settle for 10-30 min prior to picking. Fluorescent cells were picked with a mouth aspirator consisting of a 0.8 mm Nalgene Syringe Filter (Thermo), a short stretch of tubing, a plastic needle holder, and a pulled Kwik-Fil Borosilicate Glass capillary (Fisher). Cells picked off the primary plate were transferred to a Sylgard-lined 35 mm Mat Tek Glass Bottom Microwell Dishes (Mat Tek) filled with 170 microliters AHS+2%FBS, allowed to settle, and then re-picked. Three washes were performed in this way before the purified cells were picked and transferred into 3 microliters Smart-SCRB Sample Lysis buffer (0.2% Triton X-100, 0.1 U/ μL NxGen RNase Inhibitor (Biosearch Technologies) and stored at-80°C.

#### Library preparation and sequencing

For each sample, one microliter of harsh lysis buffer (50 mM Tris pH 8.0, 5 mM EDTA pH 8.0, 10 mM DTT, 1% Tween-20, 1% Triton X-100, 0.1 g/L Proteinase K (Roche), 2.5 mM dNTPs (Takara), and 1 μl 10 μM barcoded RT primer were added, and the samples were incubated for 5 m at 50°C to lyse the cells, followed by 20 m at 80°C to inactivate the Proteinase K. Reverse transcription master mix (2 μL 5X RT Buffer (Thermo-Fisher), 2 μL 5M Betaine (Sigma-Aldrich), 0.2 μL 50 μM E5V6NEXT template-switch oligo (Integrated DNA Technologies), 0.1 μL 200 U/μL Maxima H-RT (Thermo-Fisher), 0.1 μL 40U/μL NxGen RNase Inhibitor (Biosearch Technologies), and 0.6 μL nuclease-free water (Thermo-Fisher) was added to the lysis reaction and incubated at 42°C for 1.5 hr, followed by 10 min at 75°C to inactivate the reverse transcriptase. PCR was performed by adding 10 μL 2X HiFi PCR Mix (Kapa Biosystems) and 0.5 μL 60 μM SINGV6 primer and incubating at 98°C for 3 m, followed by 20 cycles of 98°C for 20 s, 64°C for 15 s, 72°C for 4 min, and a final extension of 5 min at 72°C. Samples were purified with Ampure XP Beads (0.6x ratio; Beckman Coulter), washed twice with 75% Ethanol, and eluted in 10 μL nuclease-free water. The DNA concentration of each sample was determined using Qubit High-Sensitivity DNA kit (Thermo-Fisher), and quality assessed by Bioanalyzer (Agilent).

To prepare the Illumina sequencing library, 600 pg cDNA from each pooled sample was used in a modified Nextera XT library preparation (Illumina) (Soumillon et al., 2014)using a custom set of Dual Index i5/i7 primers and extending the tagmentation time to 15 min. The resulting libraries were purified according to the Nextera XT protocol (0.6x ratio) and quantified by qPCR using Kapa Library Quantification (Kapa Biosystems). Sequencing libraries were loaded on an Illumina NextSeq High Output flow cell reading 26 bases in Read 1, including the spacer, sample barcode and UMI, 8 bases in the i7 index read, and 50 bases in Read 2 representing the cDNA fragment from the 3′ end of the transcript.

Sequencing adapters were trimmed from the reads with Cutadapt (Martin, 2011)prior to alignment with STAR (Dobin et al., 2013)to the *Drosophila* r6.17 genome assembly (Flybase). The resulting transcript alignments were passed to RSEM (Li & Dewey, 2011)to generate gene expression counts.

### 2-Photon Imaging

#### Reagents

Histamine was (Sigma Aldrich cat# 59964)was and kept in the dark at-20℃. Custom synthesized Dh44 was generated according to the previously determined peptide sequence (Cabrero et al., 2002) by Genscript. Physiological saline (in mM: 2 CaCl2, 5 KCl, 5 HEPES, 8.2 MgCl2, 108 NaCl, 4 NaHCO3, 1 NaH2PO4, 10 sucrose, and 5 trehalose; adjusted pH: ∼7.4) was prepared in advance of recordings and kept at 4℃ for no longer than 14 days. Physiological saline was warmed to room temperature prior to recording to prevent temperature interference.

#### Animal preparation

All calcium imaging experiments were performed on adult Drosophila melanogaster at ∼3-9 days post eclosion and at room temperature. Flies of the proper genotype were anesthetized on ice and then affixed to a custom built holder using UV curable glue (Bondic, cat#SK8024). For recordings from JO-A and JO-B AMMC projections the head was turned 180° so that the proboscis was pointed toward the imaging field and the antenna remained dry on the other side of the holder. Physiological saline was added to the well of the holder and then the proboscis was removed with forceps along with excess cuticle, trachea, and fat tissue. For recordings from IVLP projections of the R47F09-Gal4 AMMC B1s the head was positioned so that the ocelli and surrounding cuticle was pointed towards the imaging field and the antenna remained dry on the other side of the holder. Physiological saline was added to the well of the holder and the ocelli and surrounding cuticle as well as trachea and fat tissue was removed with forceps. All recordings were done with a constant circulation of physiological saline via a peristaltic pump (Masterflex C/L Model#77120-32).

#### Image acquisition

In vivo Ca^2+^ imaging experiments with histamine or Dh44 application were performed using a custom built (Scientifica, Clarksburg, USA) 2-photon microscope system and Mai Tai HP Ti Sapphire laser (Spectra-Physics, Milpitas, CA). Preparations were visualized with a Retiga R6 Microscope Camera (QImaging, Surrey, Canada) and a Nikon CFI LWD 16x water immersion objective. Data was acquired with a gallium arsinide phosphide (GaAsP) photomultiplier tube detector and ScanImage acquisition software (v.5.5, Vidrio Technologies). All recordings were taken at a frame rate of 6.17Hz.

In vivo Ca^2+^ imaging experiments with flight induction were performed on a custom-built two-photon microscope with an Insight DS+ pulsed laser (MKS Instruments) set at 930 nm excitation wavelength at up to 25 mW power, with emission detected through a Nikon CFI75 LWD 16X W objective by a photomultiplier tube (Hamamatsu H10770PB-40-SEL) through a 503/40 nm dichroic filter (Semrock). Volume imaging was carried out at a rate of 14.15 Hz (10 slices with 5 μm Z-step size).

#### Bath application

Histamine was added to physiological saline at different concentrations (see results) and kept from light exposure. 40μM Dh44 in saline was diluted to 1μM in physiological saline(Johnson et al., 2005). A peristaltic pump (Masterflex C/L Model#77120-32) was used to deliver saline and histamine saline or Dh44 saline to the prep at a rate of ∼1ml per minute. Drug delivery took place after the initial “saline” recording, the histamine or Dh44 saline was pumped to the prep for ∼3 minutes before beginning the recording. The sample was then washed with physiological saline for ∼5 minutes before beginning the “wash” recording.

#### Pico-injection

Glass pipettes were pulled from borosilicate glass capillary tubes (Sutter instrument cat#B100-75-10) using a glass micropipette puller (Sutter Instrument Co. model P-97). Physiological saline with 20mM histamine or 40μM Dh44 was injected into a glass pipette and delivered near the brain of each animal through a Picospritzer III (Parker, P/N 051-0600-200-003) which equated to ∼0.5μl of liquid. For each animal in these experiments the initial “saline” recording was acquired and then the animal was given a 1 minute rest. After the rest period the “pico” recording was acquired and the pico-injection took place around frame 20 of this recording and then the animal was given another 1 minute rest. After this second rest period the “170 SPI” recording was acquired and then the animal was given a 5 minute rest to allow for the saline to completely cycle from the plate. After the 5 minute rest the “580 SPI” recording was acquired. The laser shutter was closed between each recording to avoid photobleaching.

### Sound Delivery

A NI USB-6211 DAQ delivered sine waves to a servo 120a power amplifier (Samson) which transferred sound to a Koss “sparkplug” earbud attached to a tube and mounted on a micromanipulator. The tube was positioned approximately 2mm from the antenna. Custom matlab scripts defined trigger, particle velocity, frequency, length, and repetition of sound. For all recordings the particle velocity was set to 0.001 M/S, for the JO-A recordings the frequency was set to 200Hz and for the JO-B and AMMC B1 recordings the frequency was set to 100Hz. The sound stimulus was ramped up to the defined particle velocity in the first 50 ms of sound delivery with a cosine envelope, and similarly ramped down in the last 50 ms. For JO-A pico-injection experiments sound was played for 30 seconds, for all other recordings sound was played for 10 seconds. Prior to use in the above experiments, the particle velocity output of the sound system was calibrated using pure tones in the 100-1000Hz range against a custom made particle velocity microphone (Janelia, HHMI, Ashburn, VA) using custom matlab scripts. The design of the particle velocity microphone assembly was adapted from a low-noise amplifier board (two-channel version) optimized for recording *Drosophila* courtship song (Arthur et al., 2013), powering and recording from a CUI CMP-5247TF-K pressure-gradient microphone. The microphone assembly was calibrated using pure tones in the 100-1000Hz range in the acoustic far-field in an anechoic chamber against a calibrated pressure microphone using previously established methods (Arthur et al., 2010; Göpfert et al., 1999). Custom MATLAB code was used to calibrate the particle velocity microphone and sound delivery system, as well as for generation of sound stimuli.

### Flight Induction

For flight experiments while carrying out Ca^2+^ imaging in JO-As, flies were tethered via the dorsal thorax to a custom-made fly holder and dissected through the posterior side of the head. A drop of UV glue was placed on the mouthparts to minimize their impact on brain movement, and the forelegs were removed to prevent their stimulation of the antenna. For the antenna-glued experimental condition, both antennae on each animal were further fully covered in UV glue to immobilize all segments of the antenna. Flight was induced via an airpuff delivered to the fly’s anterior with a syringe attached to a 10 μL pipette tip, and flight behavior was recorded by a Blackfly S BFS-U3-04S2M-CS monochrome camera (FLIR) imaging a lateral view of the fly at 100 Hz, with image acquisition start triggered at the same time as the start of calcium imaging. Flight induction was carried out 3 times per fly.

#### ΔF/F Normalization and Peak Sound response calculation

Using ImageJ, an ROI was drawn around the neuron projections of interest and used to measure the average intensity in each frame of the recording. Another ROI was drawn in an area of the recording without any fluorescence to measure the background for each frame of the recording. The background was subtracted from the projection intensity to give the raw fluorescence intensity for each frame of the recording. The initial intensity of the projections was calculated by averaging ∼10 seconds of the recording prior to sound presentation. The initial intensity was subtracted from the raw and the result was divided by the initial to give ΔF/F. Normalization of responses was performed by dividing the ΔF/F values for each frame by the maximum ΔF/F value for the animal. Peak sound response values were calculated by averaging the ΔF/F values during each sound presentation. In JO-A bath, JO-B pico and R47F09 bath and pico samples each recording had 3 sound presentations and the average ΔF/F values for each of these sound presentations were averaged together to give the average peak ΔF/F for the recording. For JO-A pico-injection samples each recording had 2 sound presentations and the average ΔF/F values for each of these sound presentations were averaged together to give the average peak ΔF/F for the recording. For recording traces the responses were averaged with 10 seconds prior and 15 seconds post sound presentation and the SEM was calculated based on the averaged responses from each of the animals in the experiment group. For flight experiments, Z-stacks were flattened via the ImageJ sum slices Z-projection prior to ROI selection and background subtraction as described above. Baseline fluorescence was determined in the 5-second period prior to flight start and used to calculate ΔF/F for the entire trace. No further normalization was carried out.

#### Courtship behavior analysis

Flies were starved 22 hours prior to experiments. At the start of experiments, flies were cold-anesthetized prior to placement on a sloped-wall behavioral stage. Behavioral stages were custom made with white and clear acrylic plastic to fit the standard set by previous studies (Simon & Dickinson, 2010). One virgin experimental male and one virgin wild-type female were placed into each bowl. Genotypes of the males in each well were denoted with a related symbol for later analysis. The stage was placed below a behavioral camera (FLIR Flea 2.0) shooting 40 fps and above a custom made IR light array, made with 910nm IR LEDs, for optimum visualization of appendages(Scaplen et al., 2019). After waiting 15 minutes to allow flies to wake up and acclimate, a 10 minute recording was made of the fly behavior, based on the average time to copulate. Analysis of videos was performed to quantify courtship progress and copulatory success. As courtship is hierarchical, progressive steps through male courtship were scored on an ordinal scale: 0=no courtship attempts; 1=orientation; 2=tap; 3=wing extension/singing; 4=sniff/lick; 5=mount; 6=successful copulation (Spieth, 1974). For all recordings, each attempt to court the female was scored independently.

#### Statistical analysis

All statistical analyses were performed using the software GraphPad Prism Version 10. All data was first tested for normal distribution using the D’Agostino & Pearson normality test. For average peak sound response comparisons between males and females two way repeated measures ANOVA tests were performed with values matched per animal and post-hoc Tukey’s test for individual comparisons. For JON receptor expression, AMMC B1 Dh44 receptor expression in males and females, average peak sound responses without sex comparison, and copulation success proportions one-way ANOVA tests were performed with post-hoc Tukey’s test for individual comparison and values were paired where applicable. For RNAi validation, JO-A activity during flight, and histamine fluorescence quantification in the AMMC unpaired t-tests were performed. Significance values are indicated as follows: P>0.05 = NS, P<0.05 = *, P<0.01 = **, P<0.001 = ***, P<0.0001 = ****.

## Supplementary Figures

**Figure S1.1.**
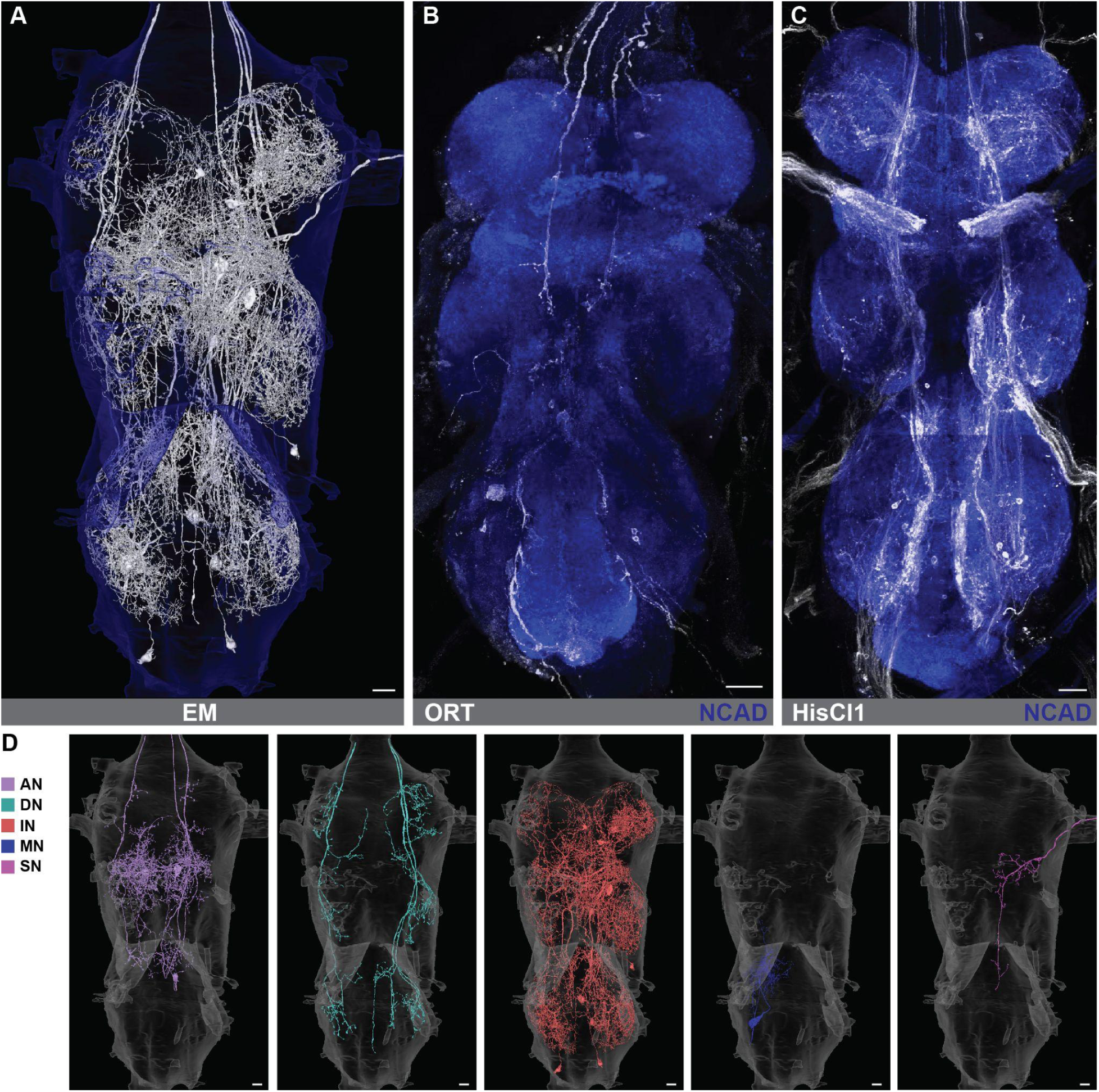
Top VNC targets of the MtAHNs do not match histamine receptor Gal4 Expression. **(A)** Top downstream targets of the MtAHNs in the VNC EM dataset (MANC). **(B)** TI{2A-Gal4}ort[2A-Gal4] driving expression of GFP (white) in the VNC with NCAD as a neuropil marker (blue). **(C)** T2A-Gal4{HisClB}T2A-Gal4 driving expression of GFP (white) in the VNC with NCAD as a neuropil marker (blue). **(D)** Top downstream targets of the MtAHNs in the VNC with colors corresponding to cell class as indicated by the legend on the left: ascending neuron (AN;purple), descending neuron (DN;cyan), interneuron (IN;red), motor neuron (MN;blue), and sensory neuron (SN;magenta). Scale bar = 30μm.

**Figure S1.2.**
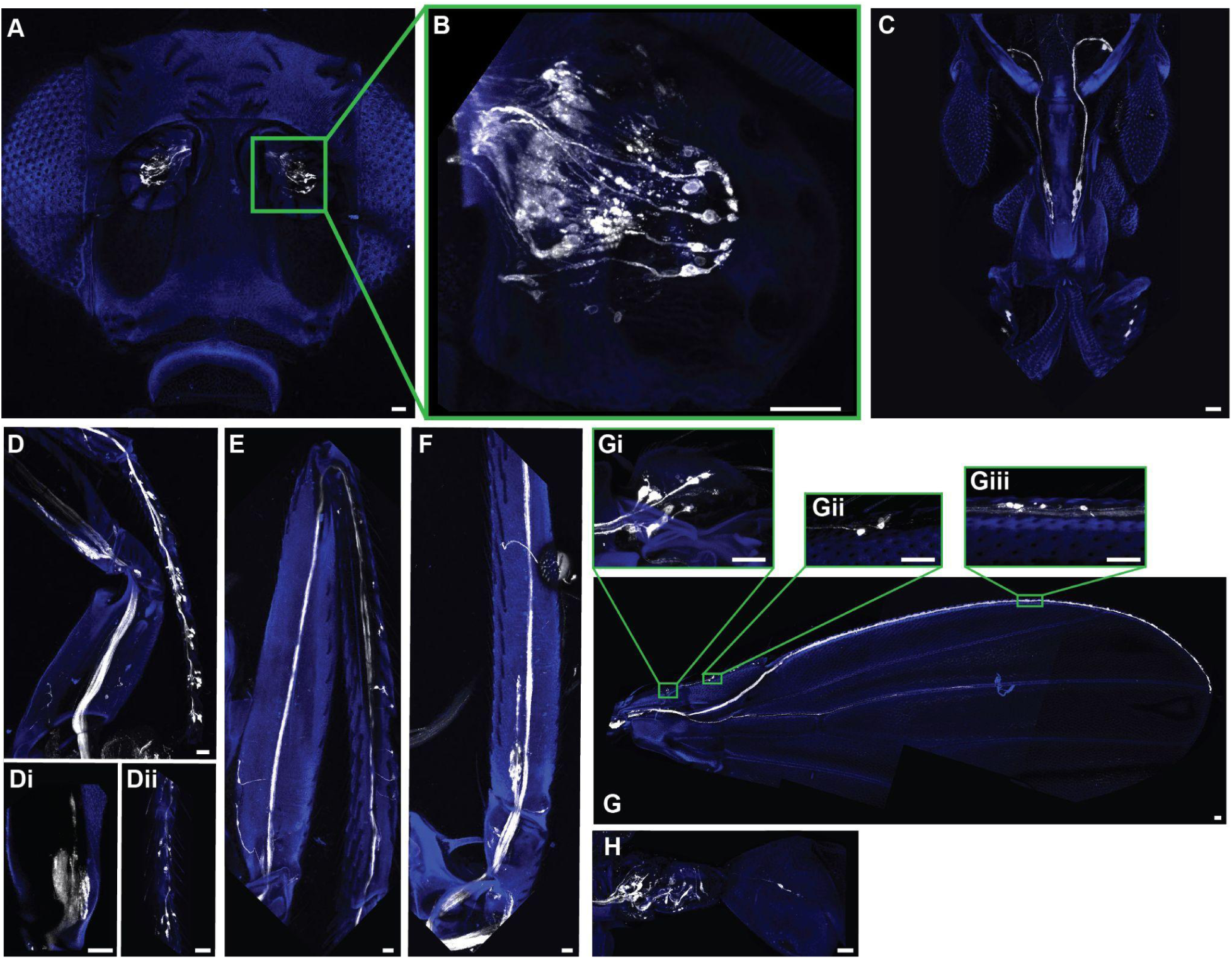
HisCl1-Gal4 expression in appendages. HisCl1-Gal4 driven expression of GFP (White) in Johnston’s organ **(A,B)**, proboscis **(C)**, front leg **(D)** including femoral chordotonal organ**(Di)** and tarsi **(Dii)**, hind leg **(E)**, mid leg **(F)**, wing **(Gi-iii)**, and haltere **(H)** with cuticle autofluorescence (Blue). All scale bars = 20 μm.

**Figure S1.3.**
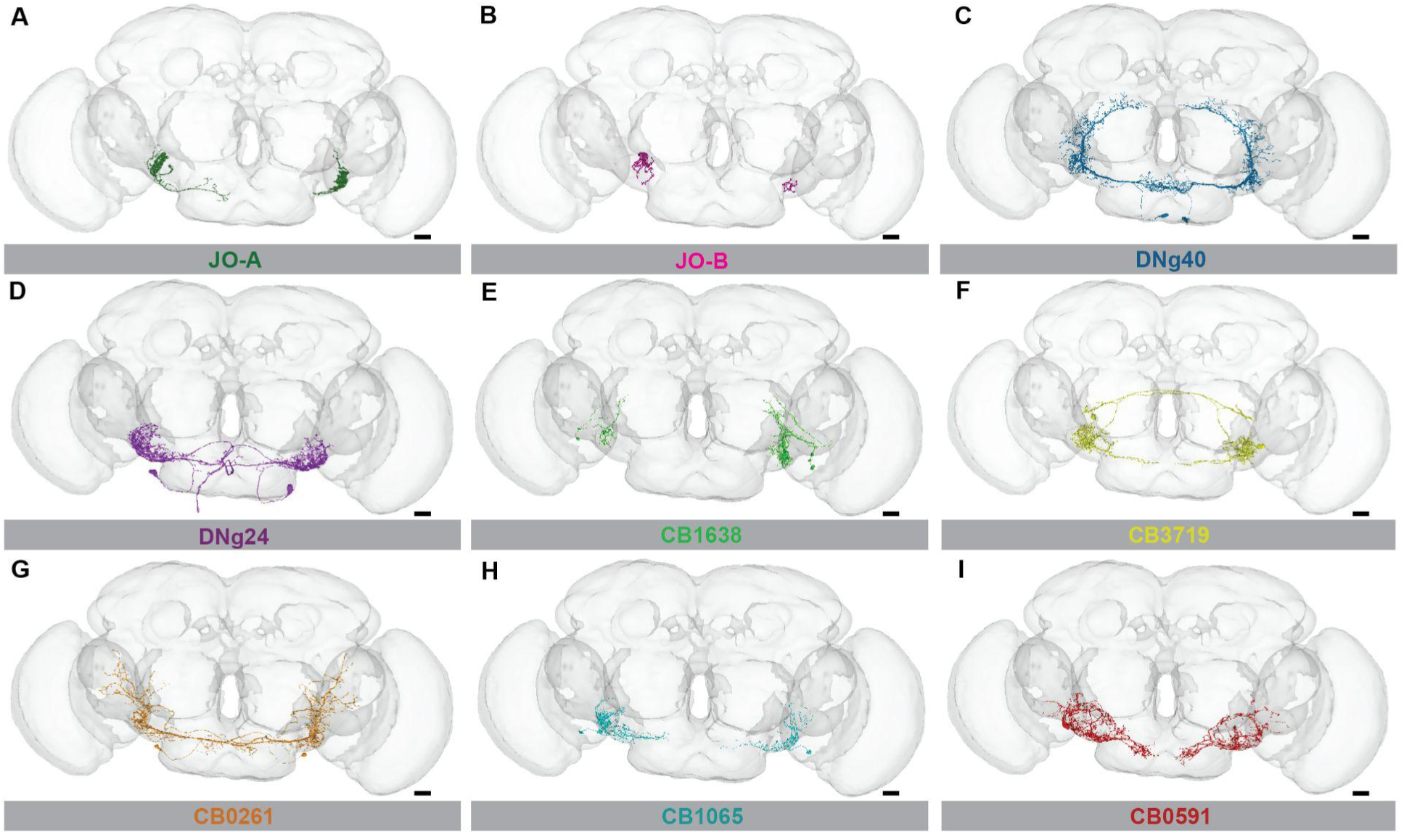
Top downstream targets of MtAHNs in AMMC split by cell type. (A-I) Top cell types downstream of MtAHNs in the AMMC from the brain EM dataset (FAFB), cell type as annotated in Flywire, indicated below each image. All scale bars = 20 μm.

**Figure S1.4.**
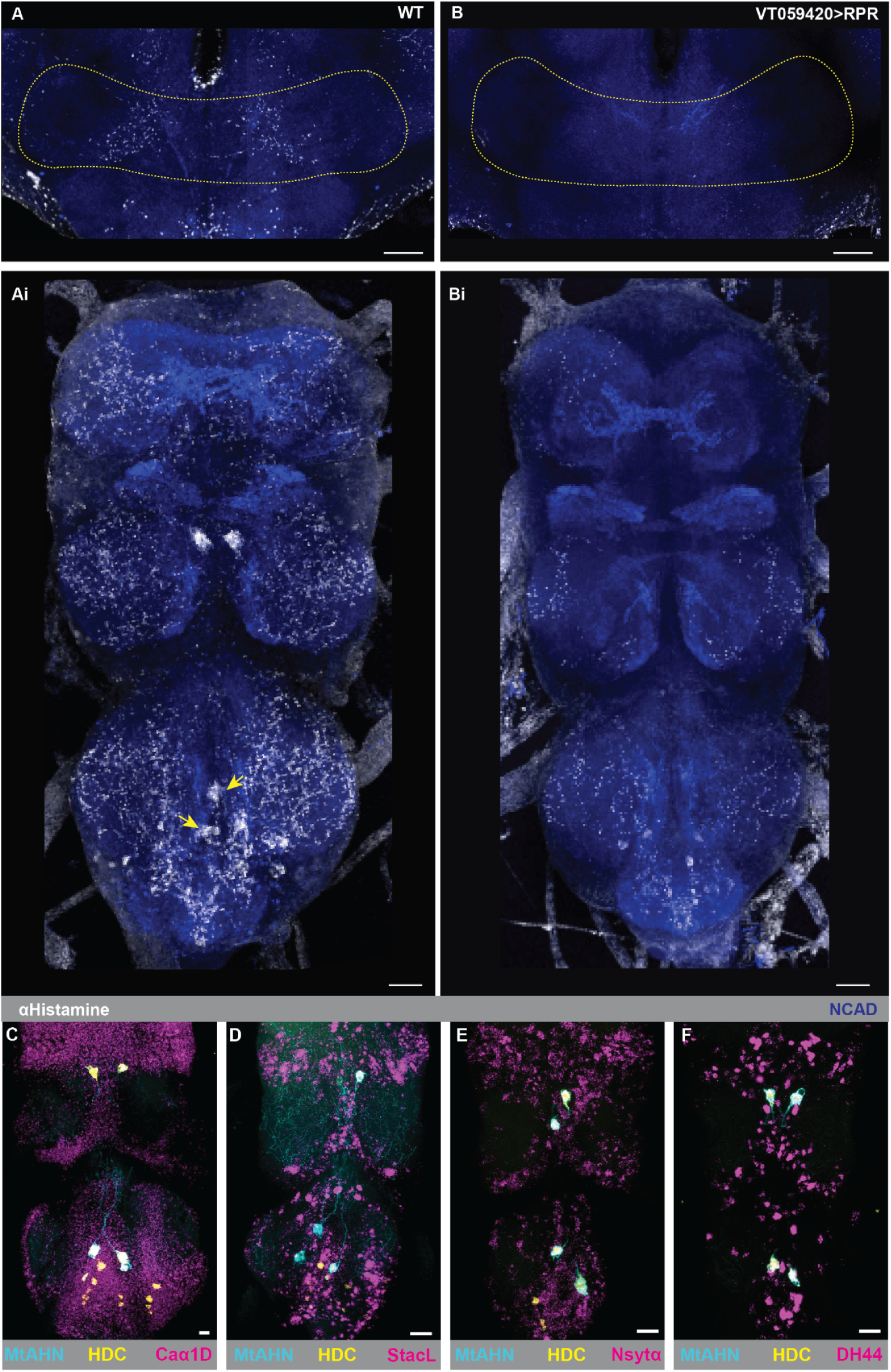
Confirmation of MtAHN histaminergic output regions and HCR labeling in VNC. (A-B) Histamine antibody labeling (white) in the AMMC indicated with yellow dotted line with NCAD as a neuropil marker (blue) in **(A)** WT and **(B)**;;VT059420-LexAGADfl>Aop-RPR flies. **(Ai-Bi)** Histamine antibody labeling (white) in the VNC with NCAD as a neuropil marker (blue) and MtAHN cell bodies indicated with yellow arrows in **(Ai)** WT and **(Bi)**;;VT059420-LexAGADfl>Aop-RPR flies. **(A-Bi)** Scale bars = 30μm. **(C-F)** GFP expression (Cyan) driven by VT025938AD ∩ VT040583DBD-splitGal4 with HDC HCR-labeling (Yellow) as verification of histaminergic cells and **(C)** Caα1D (Magenta), **(D)** StacL (Magenta), **(E)** Nsytα (Magenta), **(F)** Dh44 (Magenta) expression in the VNC **(C-F)** scale bars = 20μm.

**Figure S1.5.**
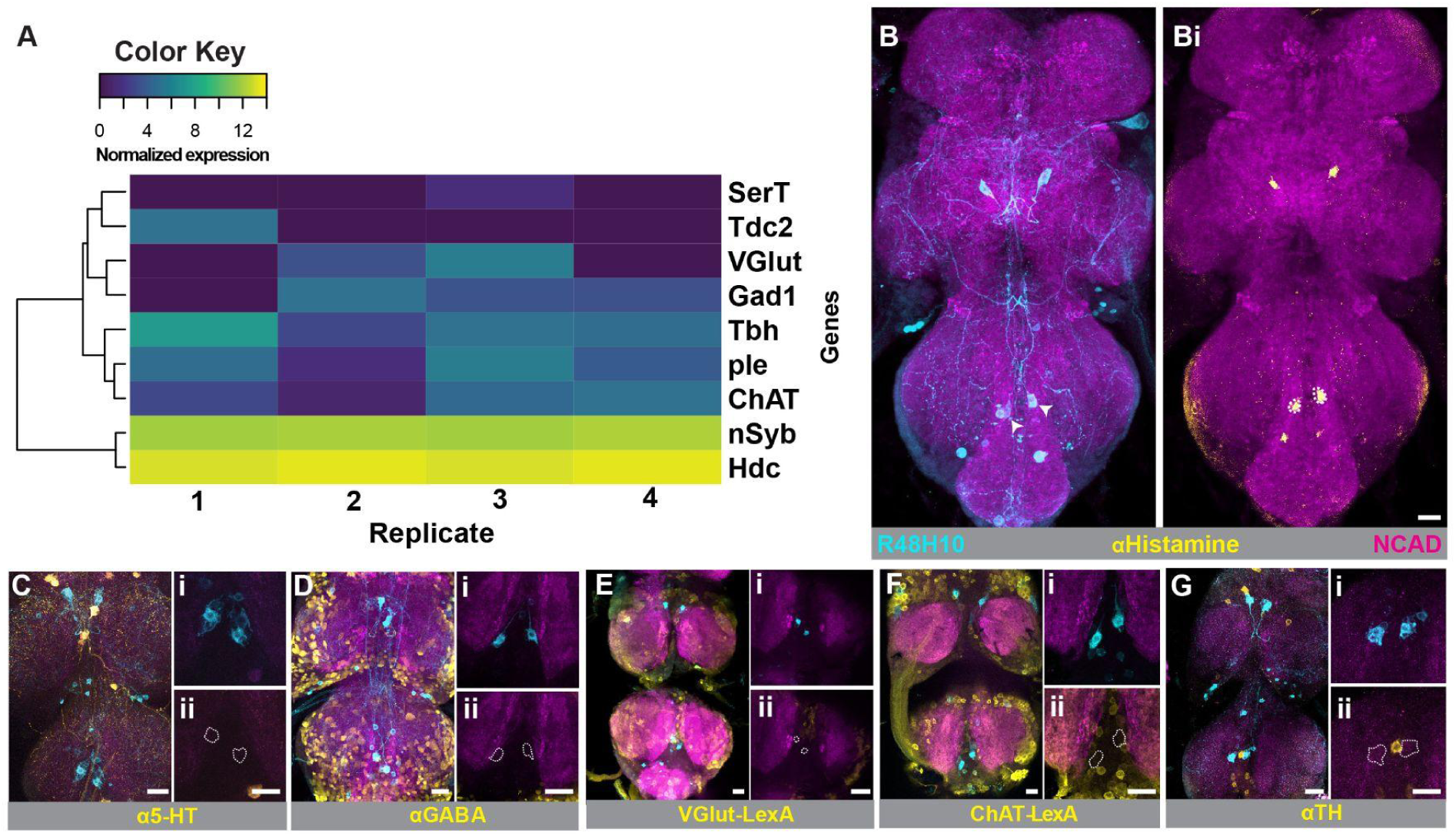
MtAHN RNAseq results for canonical neurotransmitters compared to immunolabeling. **(A)** Normalized expression of rate limiting enzymes or vesicular transporters for small classical transmitters with Nsyb as a neuronal cell marker from MtAHN RNAseq. Log10 normalized expression level indicated with color key. **(B-D,G)** R48H10-Gal4 driven expression of GFP (cyan) with NCAD (magenta) as a neuropil marker. **(B)** MtAHN cell bodies indicated with white arrows and **(Bi)** white dotted lines in histamine signal (yellow) channel. **(C)** Serotonin (5-HT) expression immunolabeling in VNC (yellow) **(Ci-ii)** with MtAHN cell body location indicated with white dotted lines. **(D)** GABA expression immunolabeling in VNC (yellow) **(Di-ii)** with MtAHNs cell body location indicated with white dotted lines. **(E)** R48H10-Gal4 driving expression of RFP (Cyan) with VGlut-LexA driving GFP expression in VNC (yellow) **(Ci-ii)** with MtAHNs cell body location indicated with white dotted lines. **(F)** R48H10-Gal4 driving expression of RFP (Cyan) with CHAT-LexA driving GFP expression in VNC (yellow) **(Fi-ii)** with MtAHNs cell body location indicated with white dotted lines. **(G)** TH expression immunolabeling in VNC (yellow) **(Gi-ii)** with MtAHNs cell body location indicated with white dotted lines. All scale bars = 10 μm.

**Figure S1.6.**
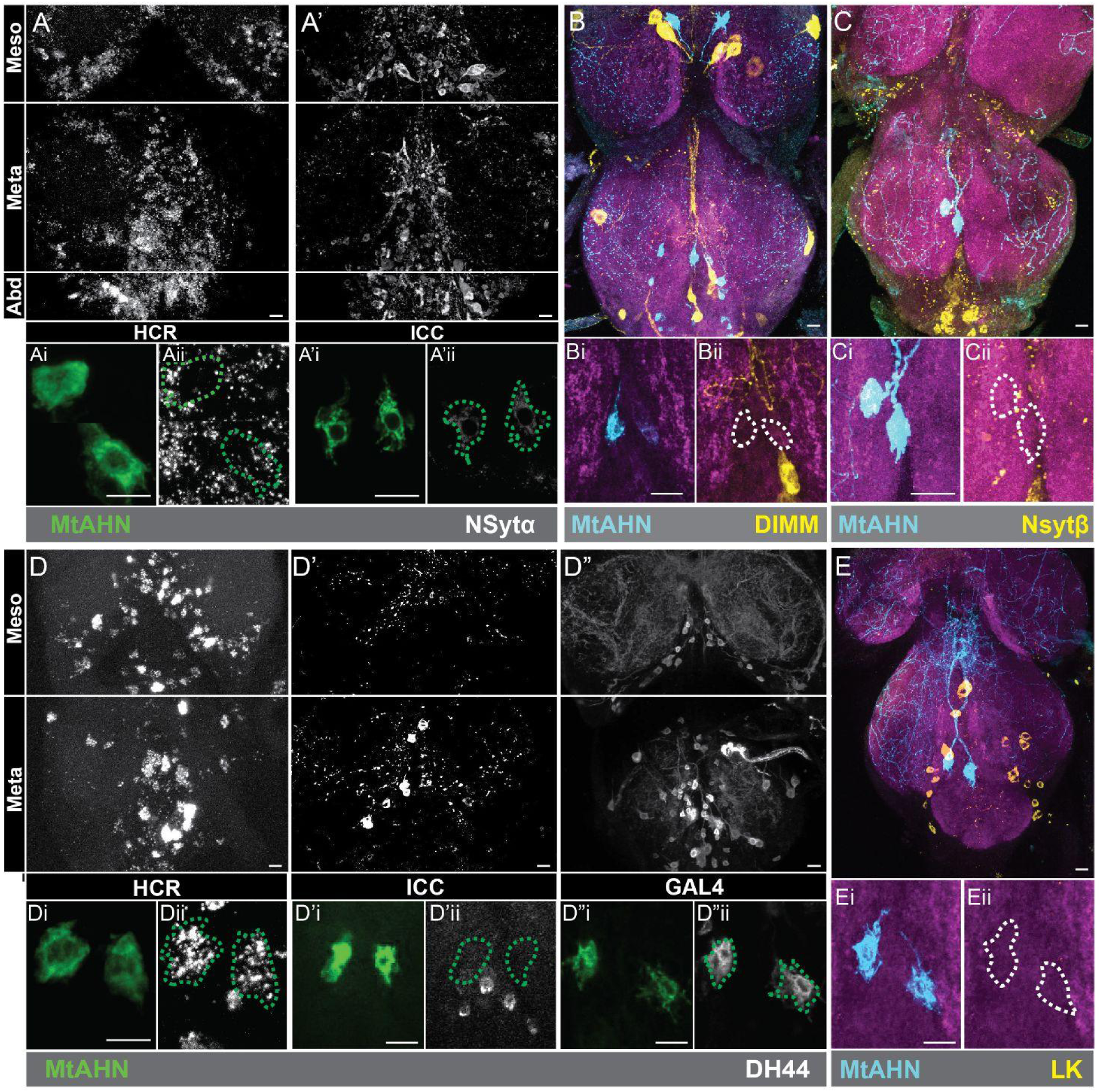
HCR and immunolabeling indicate the MtAHNs express Dh44 and Nsytα but not leucokinin, DIMM, or Nsytβ. (A-A’) NSytα HCR compared to antibody labeling (white) in the metathoracic neuromere with cell body single optical section images **(Ai-ii, A’i-ii)** of the MtAHNs (green) and cell body location indicated with green dotted lines. **(B)** Histamine antibody labeling (cyan) in the VNC with GFP expression (yellow) driven by P{w[+mW.hs]=GawB}crc[929] **(Bi-ii)** with cell body images of the MtAHNs and their location indicated with white dotted lines. **(C)** GFP expression (cyan) driven by the VT025938AD∩VT040583DBD splitGal4 with NSytβ antibody labeling (yellow) in the VNC **(Ci-ii)** with cell body images of the MtAHNs and their location indicated with white dotted lines. **(D-D”)** Dh44 HCR labeling compared to Dh44 antibody labeling and TI{2A-Gal4}Dh44[2A-Gal4] driven expression of GFP (white). **(Di-ii, D’i-ii)** MtAHN cell body, single optical section, images of VT025938AD∩VT040583DBD splitGal4 driven expression of GFP (green) with their location indicated with green dotted lines. **(D”i-ii)** MtAHN cell body images of histamine antibody labeling (green) with their location indicated with green dotted lines. **(E)** Leucokinin (LK) antibody labeling in the VNC (yellow) with GFP expression driven by VT025938AD∩VT040583DBD splitGal4 (cyan) and **(Ei-ii)** cell body images of MtAHNs with their location indicated with white dotted lines. All scale bars = 10 μm.

**Figure S1.7.**
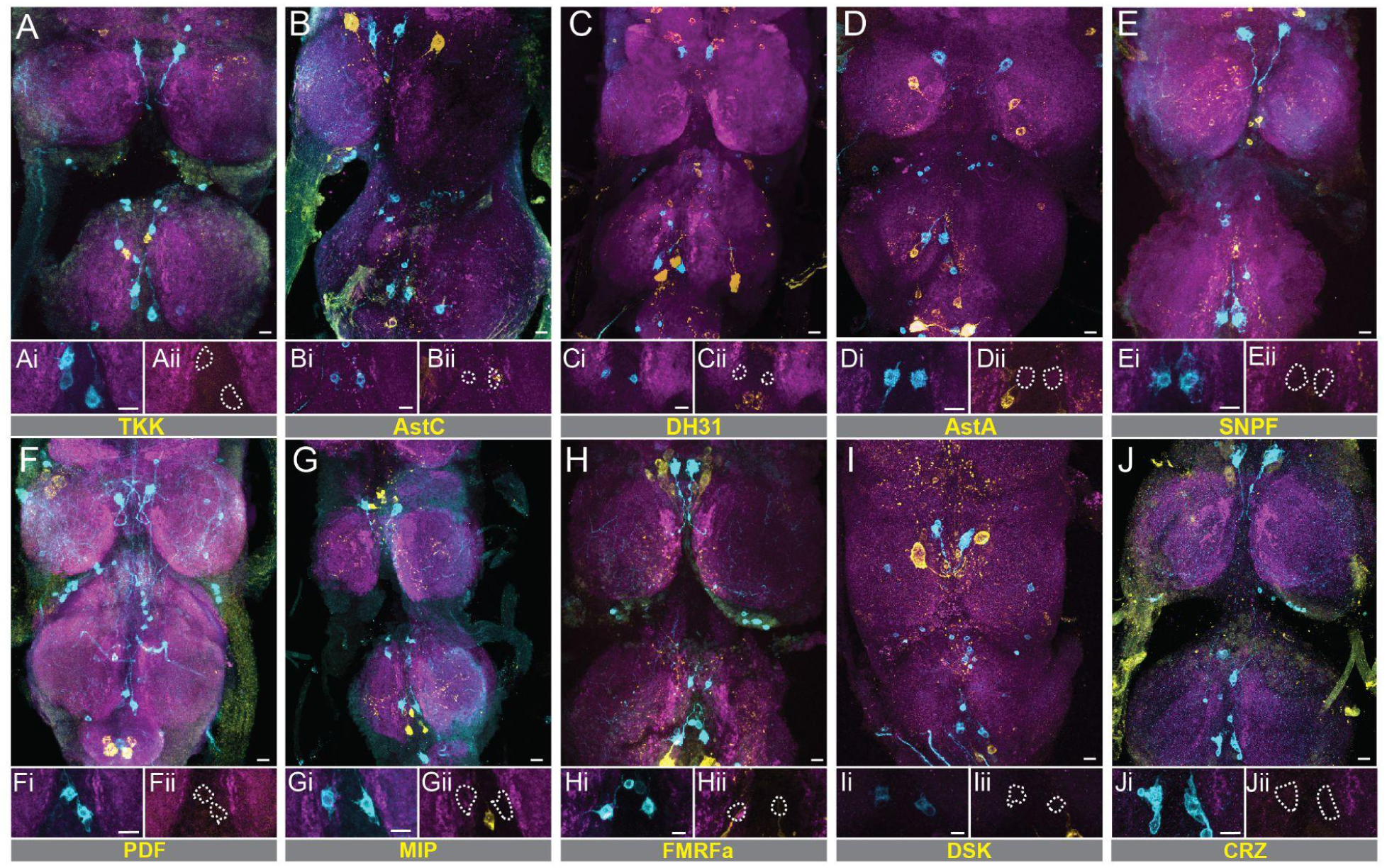
Neuropeptide antibody labeling shows no overlap with the MtAHN cell bodies. (A-J) GFP expression (cyan) driven by R48H10-Gal4 with **(A)** Tachykinin (TKK), **(B)** Allatostatin C (AstC), **(C)** DH31, **(D)** Allatostatin A (AstA), **(E)** SNPF, **(F)** PDF, **(G)** MIP, **(H)** FMRFamide (FMRFa), **(I)** Drosulfakinin (DSK), and **(J)** Corazonin antibody labeling (yellow). **(Ai-Ji)** MtAHN cell body images **(Aii-Jii)** with neuropeptide antibody labeling and white dotted lines indicating MtAHN cell body location. All scale bars = 10 μm.

**Figure S2.1.**
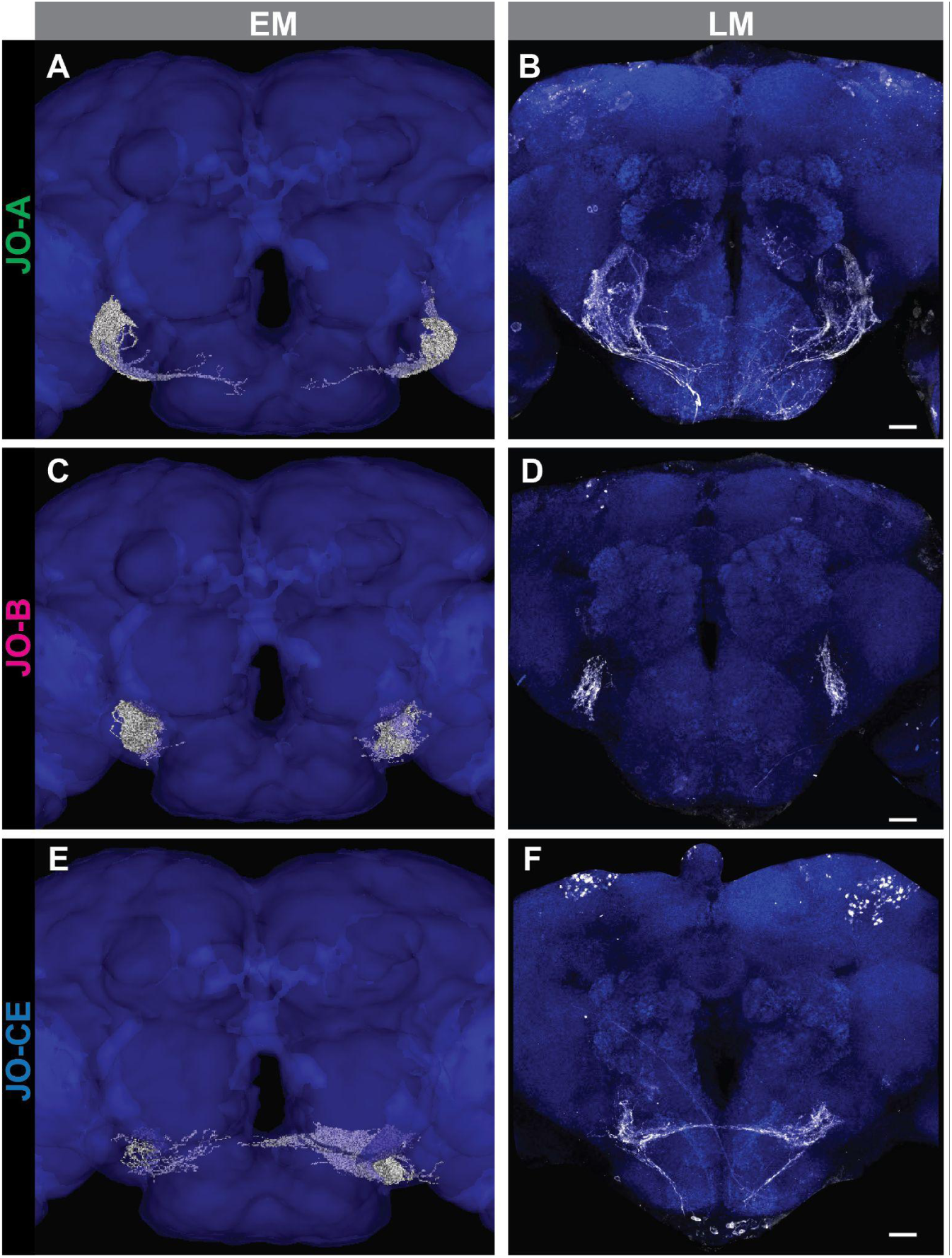
Comparison of EM reconstruction to LM images of JON subtypes in the brain. (A,C,E) JONs in the FAFB dataset annotated as **(A)** JO-A, **(C)** JO-B, **(E)** JO-C, JO-D, and JO-E (white) overlaid on neuropil mesh (blue). **(B,D,F)** GFP expression (white) driven by **(B)** R74C10-Gal4, **(D)** NP1046-Gal4, and **(E)** NP6250-Gal4 with NCAD (blue) as a neuropil marker. All scale bars = 20 μm.

**Figure S3.1.**
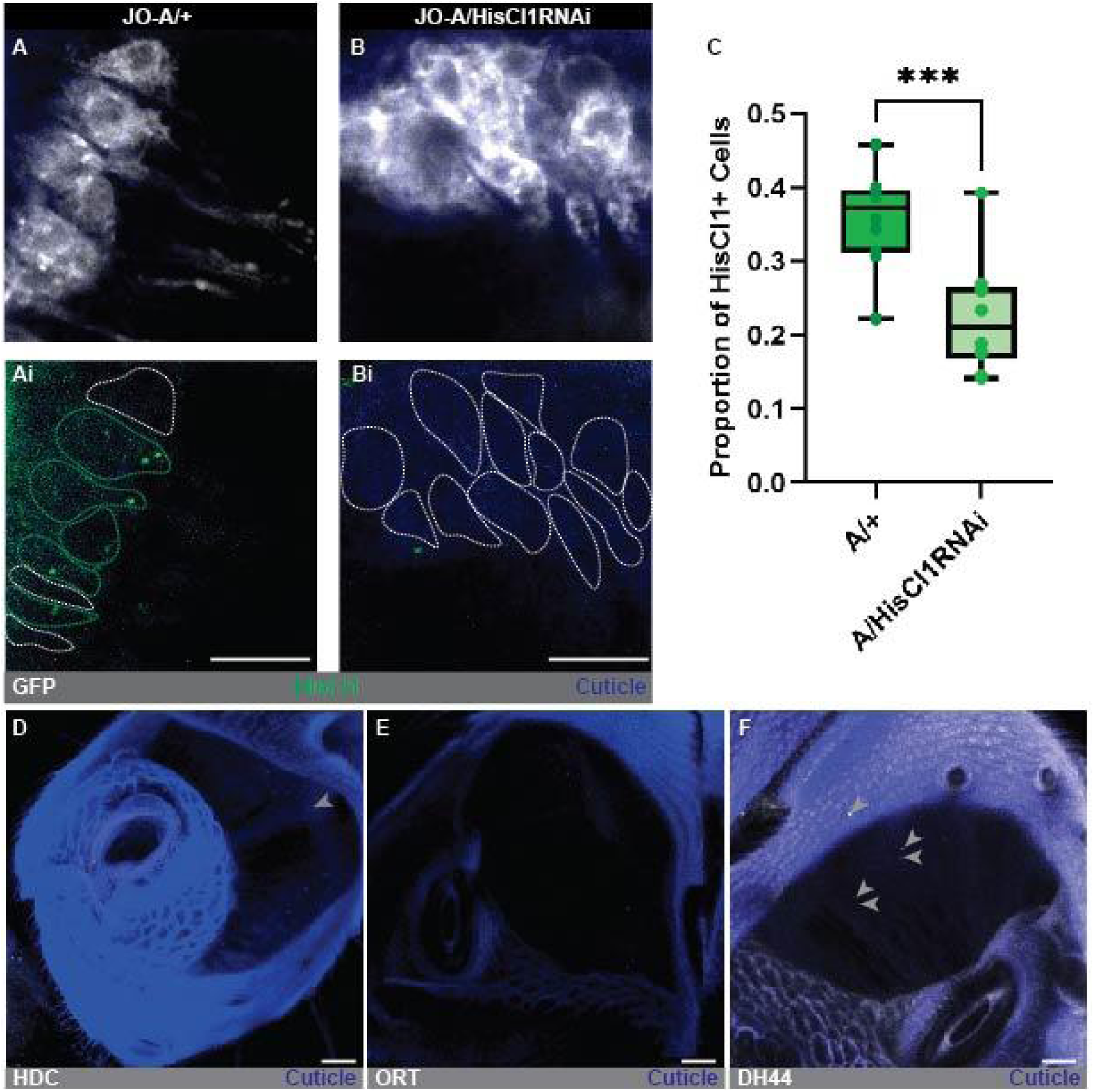
HisCl1 RNAi confirmation and expression of HisCl1, Ort, HDC, and Dh44 in the JO. (A-B) Single optical section image of GFP expression driven by R74C10-Gal4 (white) with **(B)** HisCl1 expression knockdown mediated by RNAi. **(Ai-Bi)** Cell body images of JO-As with HisCl1 HCR labeling (green), signal overlap indicated with green dotted lines and non-overlap indicated with white dotted lines. **(C)** Box-plot of HisCl1 expression quantification in heterozygous samples of R74C10-Gal4 and R74C10-Gal4>UAS-HisCl1-RNAi with significance indicated above bars (N=10 antenna, significance from unpaired t-test: ***=P<0.001, Error bars = STDEV). **(D)** Max projection of histidine decarboxylase (HDC) HCR labeling (white) in JO with cuticle autofluorescence (blue)and puncta indicated with grey arrows. **(E)** Max projection of ORT HCR labeling (white) in JO with cuticle autofluorescence (blue). **(F)** Max projection of Dh44 HCR labeling (white) in JO with cuticle autofluorescence (blue). All scale bars = 20μm.

**Figure S3.2.**
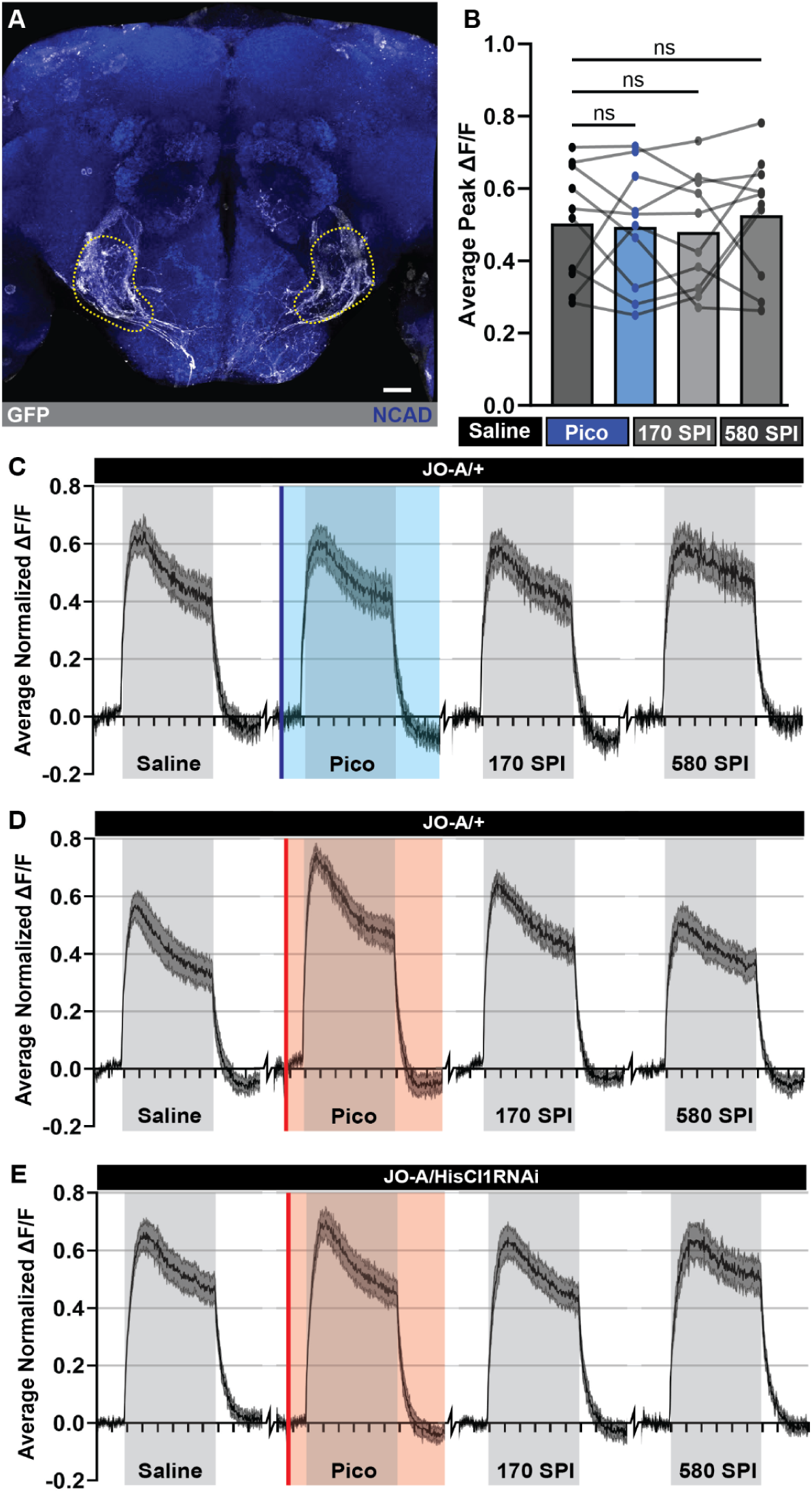
Saline controls and averaged traces for JO-A neuron recordings with pico-injections. **(A)** R74C10-Gal4 driven expression of GFP (white) in the brain with NCAD (blue) as a neuropil marker, JO-A AMMC projections indicated with yellow dotted lines scale bar set to 20μm. **(B)** Comparison of R74C10>GCaMP7f responses between initial saline recording, saline pico injection, 170 seconds post-injection (SPI), and 580 SPI significance indicated above bars (N=10, significance from repeated measures ANOVA with post-hoc Tukey’s test for individual comparisons: NS=P>0.05). **(C-E)** Recording traces of heterozygous R74C10>GCaMP7f (N=10, average ΔF/F ± SEM) responses to sound (grey) with (C) saline pico injection (blue, solid line: injection, box: length of recording), (D,E) histamine pico-injection (red, line: injection, box: length of recording), and (E) HisCl1 expression knockdown mediated by HisCl-1RNAi, X axis is time in 5 second intervals.

**Figure S3.3.**
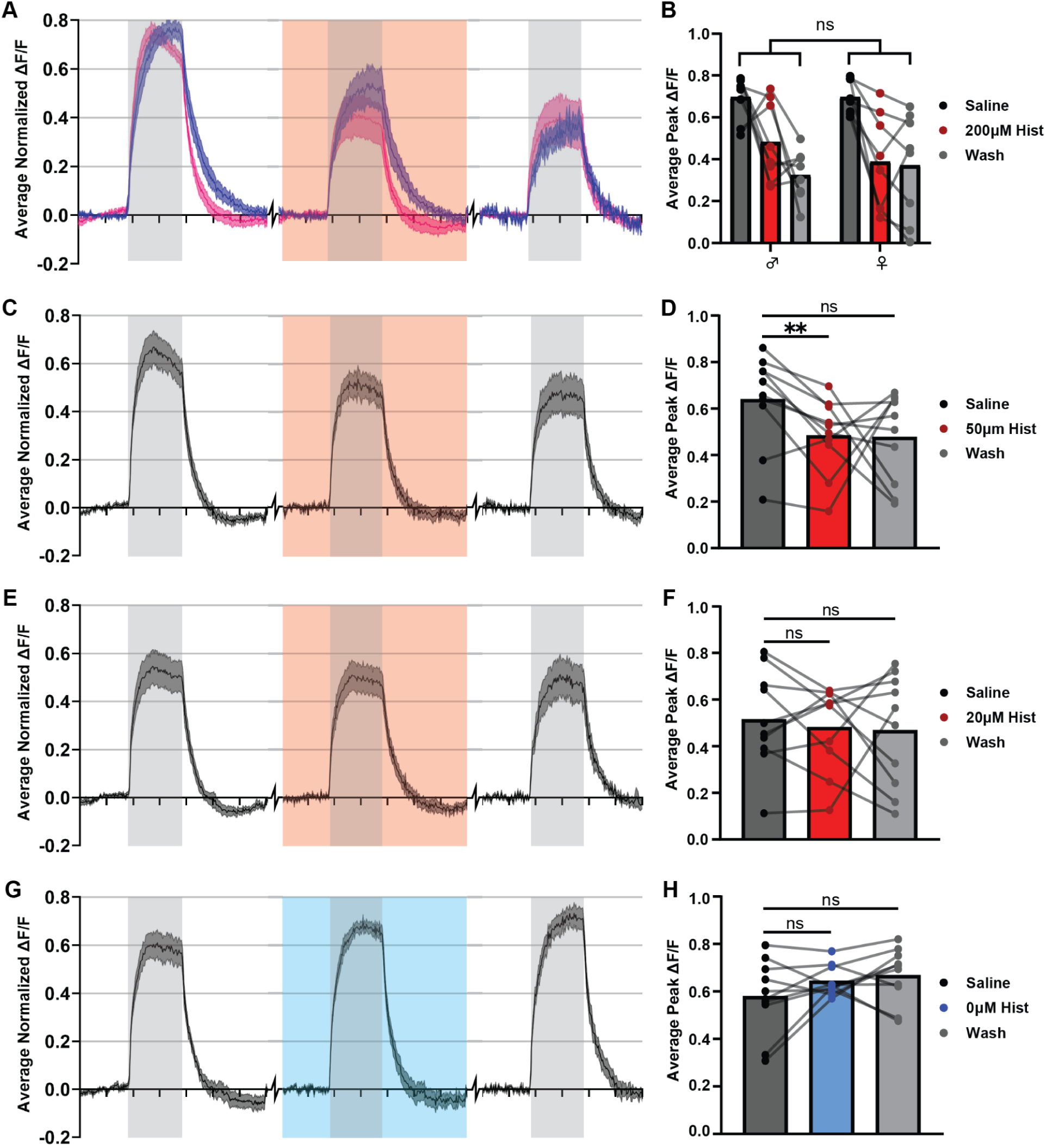
Histamine bath application with JO-A>GCaMP7f sound response traces. (A, C, E, **G)** Ca^2+^ traces of R74C10>GCaMP7f flies (N=10, average ΔF/F ± SEM) exposed to sound (grey) with addition of histamine (red) at concentrations of **(A)** 200μM, **(C)** 50μM, and **(E)** 20μM, or **(G)** continued saline (blue). **(A)** Comparison of male (blue) and female (pink) sound responses with bath application of 200μM histamine. **(B, D, F, H)** Before and after plots of average ΔF/F during sound exposure under initial saline conditions (black), **(B, D, F)** histamine bath application (red) or **(H)** continued saline (blue), and post bath application wash (grey) with histamine concentration indicated in legend to the right (N=10, significance from repeated measures ANOVA with post-hoc Tukey’s test for individual comparisons: NS=P>0.05, **=P<0.01). **(Bi)** Before and after plots of average ΔF/F during sound exposure in male and female flies (indicated on X-axis) (N=10, significance from 2-way repeated measures ANOVA: NS=P>0.05).

**Figure S3.4.**
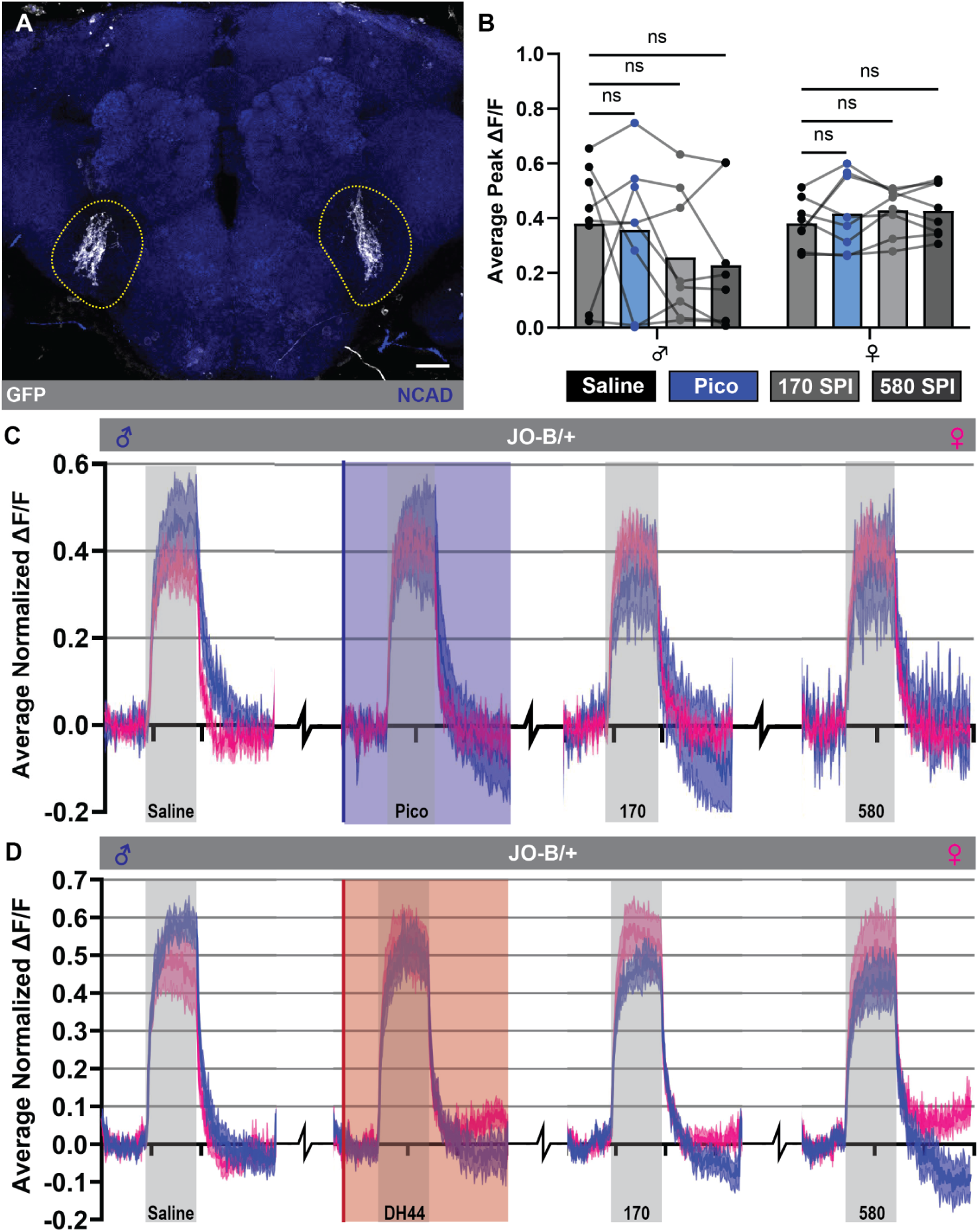
JO-B pico injection control and Dh44 application Ca^2+^ traces. **(A)** NP1046-Gal4 driving expression of GFP (White) in JO-B AMMC projections (yellow dotted line) with NCAD (blue) as a neuropil marker. **(B)** Before and after plot of average ΔF/F during sound presentation in JO-B AMMC projections from NP1046>GCaMP7f male and female flies (indicated on x-axis) under initial saline conditions (black), saline pico injection (blue), 170 SPI (light grey), and 580 SPI (dark grey) (N=13, significance from 2-way repeated measures ANOVA with post-hoc individual comparisons: NS=P>0.05). **(C-D)** Ca2+ traces (average ΔF/F ± SEM) from JO-B AMMC projections in NP1046>GCaMP7f flies with sound presentation (grey) and **(C)** saline pico injection (N=13, blue, line: point of injection, box: length of recording) or **(D)** Dh44 pico injection (N=8, red, line: point of injection, box: length of recording) and sexes indicated with separate colors (Male=blue, female=pink) (X-axis=time in 5 second intervals).

**Figure S3.5.**
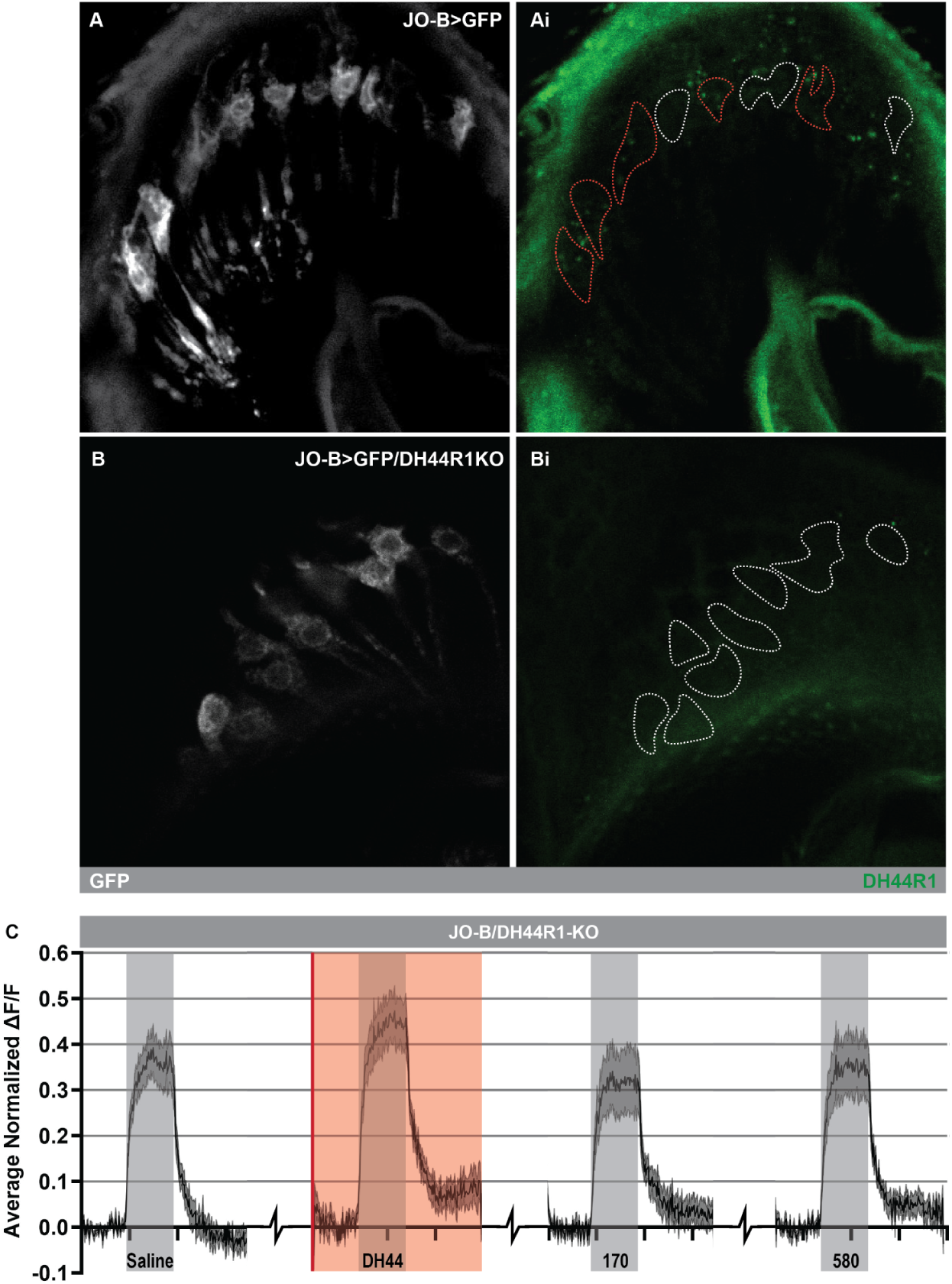
Crispr mediated knock-out of Dh44R1 in the male JO-Bs reduces HCR labeling and eliminates the sound response suppression from Dh44 application. (A-B) Single optical sections of male JO-B cell bodies (white) of NP1046-Gal4 driving expression of GFP and **(B)** crispr mediated knockout of Dh44R1. (Ai-Bi) HCR labeling for Dh44R1 (green) with JO-B cell body location indicated with dotted lines, red indicates Dh44R1 positive cells. **(C)** Ca2+ traces (average ΔF/F ± SEM) from male JO-B AMMC projections in NP1046>GCaMP7f/Dh44R1-KO flies with sound presentation (grey) and Dh44 pico injection (N=10,point of injection: red line, recording length: red box, x-axis=time in 5s intervals)

**Figure S4.1.**
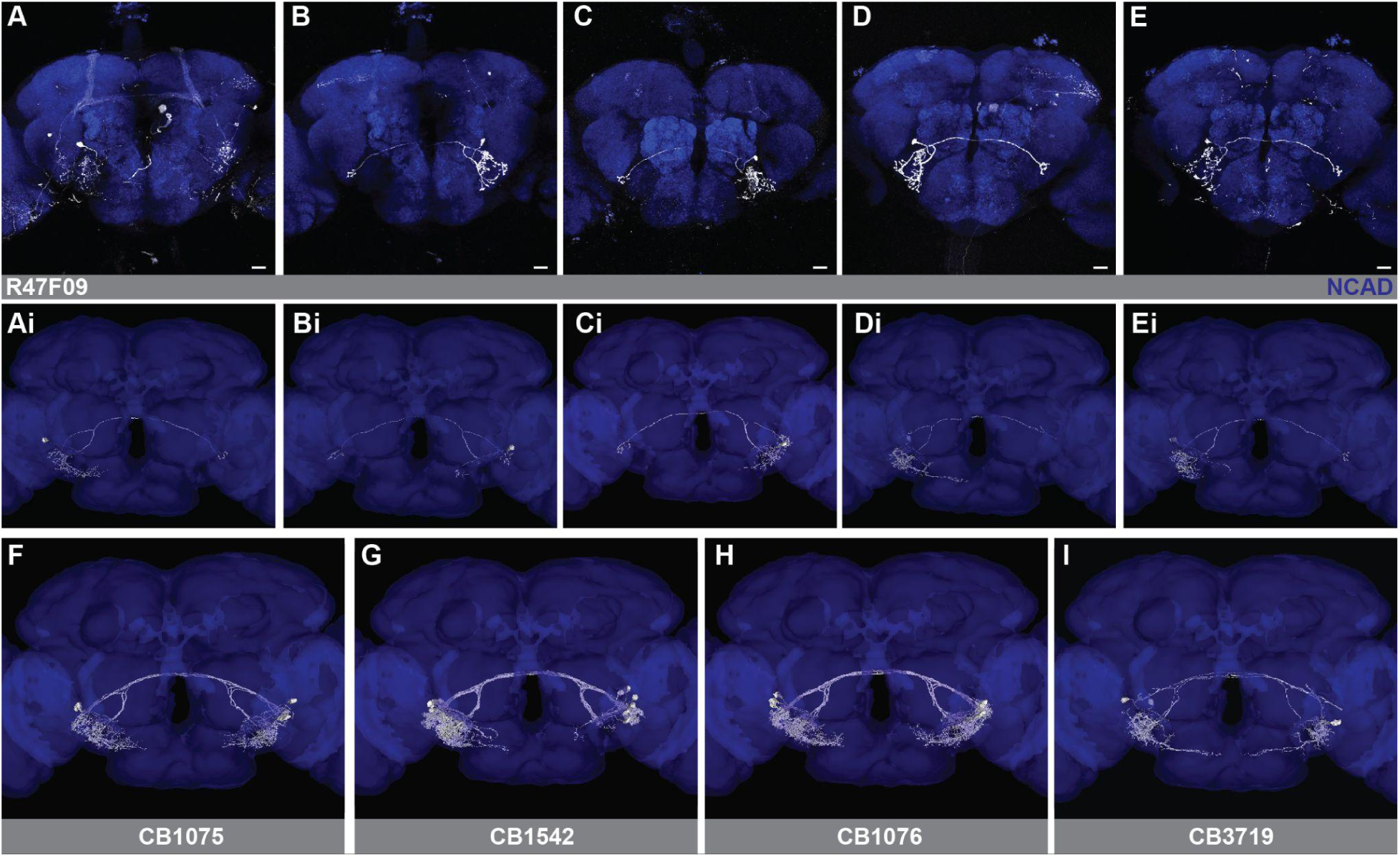
Neuronbridge search results for R47F09-Gal4 AMMC B1 neurons. (A-E) Stochastic labeling of neurons in the R47F09hGal4 (white) line driven by MCFO with NCAD antibody labeling of neuropil (blue). Scale bars = 10μm. **(Ai-Ei)** Top neuron match from neuronbridge search for each of the above images. **(F-I)** All neurons in each cell type from the neuronbridge match according to the FlyWire annotations.

**Figure S4.2.**
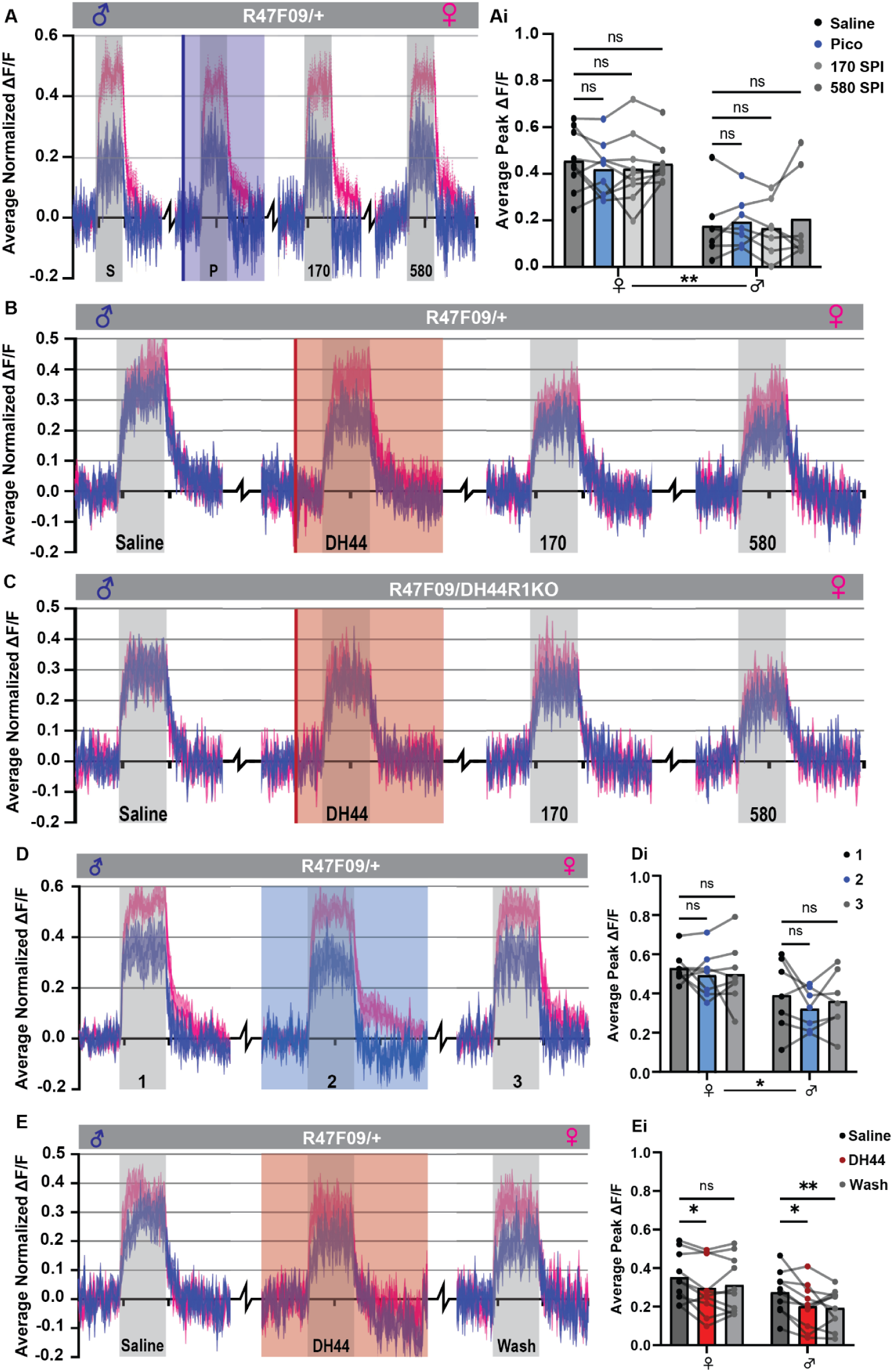
Pico-injection and bath application of Dh44 decreases sound responses in the R47F09 AMMC B1s. (A-E) Ca2+ recording traces (average ΔF/F ± SEM) from IVLP projections of AMMC B1s in the R47F09-Gal4 driving expression of GCaMP7f from male (blue) and female (pink) flies, sound presentation indicated with grey boxes (X-axis=time in 10 second intervals). **(A)** Pico-injection of saline (point of injection: blue line, length of recording: blue box). **(Ai)** Before and after plots comparing AMMC-B1 sound responses during initial saline (black), saline pico-injection (blue), 170 SPI (light grey), and 580 SPI (dark grey) in male (N=7) and female flies (N=9) as indicated on x-axis (significance from repeated measures ANOVA with post-hoc individual comparisons (above): NS=P>0.05, significance from 2-way repeated measures ANOVA (below): **=P<0.01). **(B-C)** Dh44 pico injection (red line: point of injection, red box: length of recording) **(C)** with CRISPR mediated Dh44R1-KO. **(D-E)** AMMC-B1 sound responses during constant saline bath (blue box) or Dh44 bath application (red box). **(Di-Ei)** Before and after plots indicating average ΔF/F during sound presentation with dots representing each animal and lines connecting paired values, colors correspond to recording conditions indicated in the legend (left) (significance from repeated measures ANOVA with post-hoc Tukey’s test for individual comparisons (above): NS=P>0.05, *=P<0.05, **=P<0.01) (significance from two way repeated measure ANOVA (below): *=P<0.05) **(Di)** N=8, **(Ei)** N=10.

**Figure S4.3.**
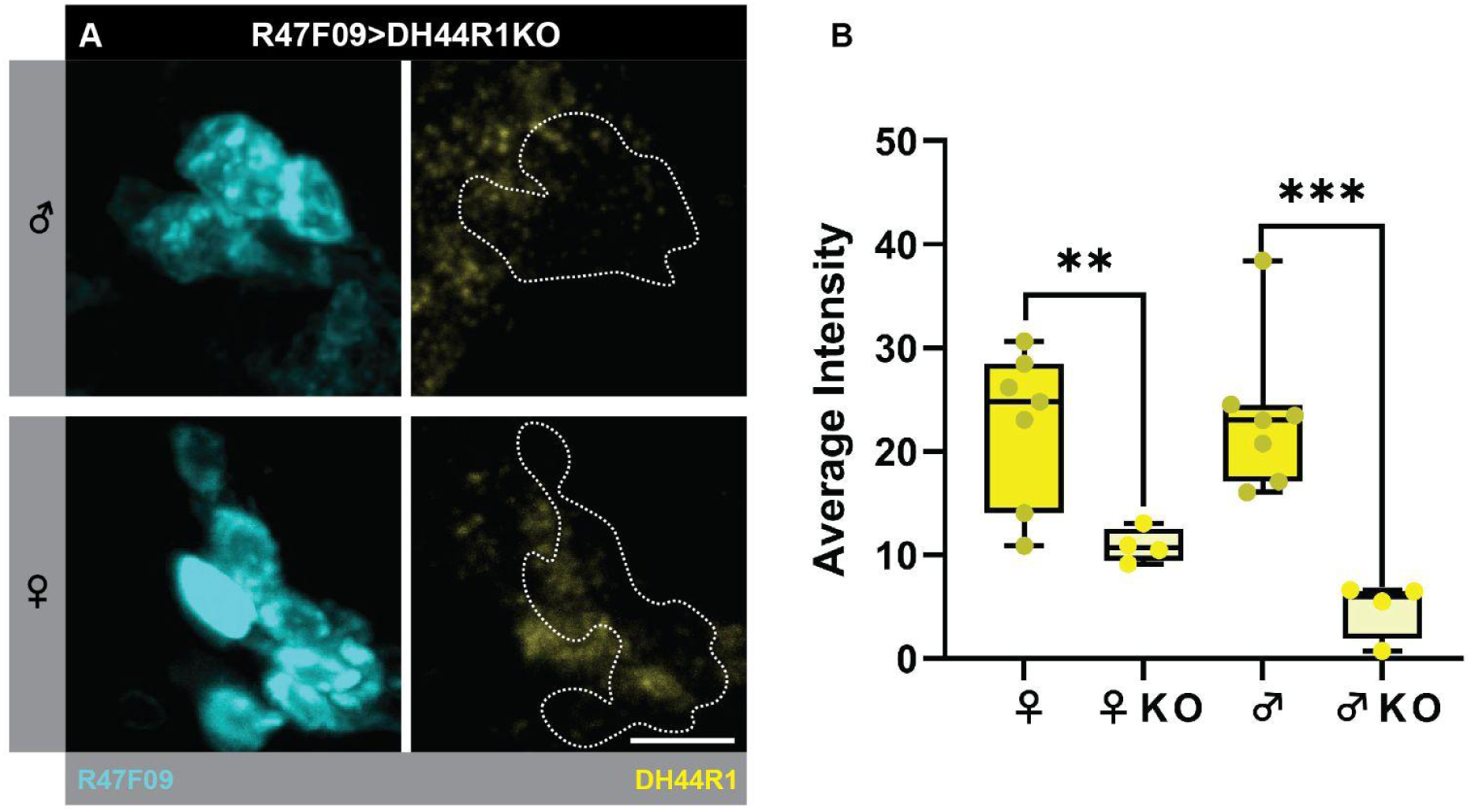
Confirmation of Dh44R1 knock out in R47F09-Gal4 AMMC B1 neurons. **(A)** Cell body images from male and female (indicated left) AMMC B1 neurons with GFP (cyan) driven by R47F09-Gal4 and Dh44R1 HCR labeling (yellow) with cell body location indicated via white dotted lines (scale bar=10μm). **(B)** Boxplot of Dh44R1 HCR labeling intensity from single optical slices of male and female AMMC B1 neurons in R47F09-Gal4 (N=7) compared to Dh44R1 knock out samples (N=4) (sex and genotype indicated on X-axis, significance from unpaired t-tests: **=P<0.01, ***=P<0.001, error bars=STDEV)

**Figure S5.1.**
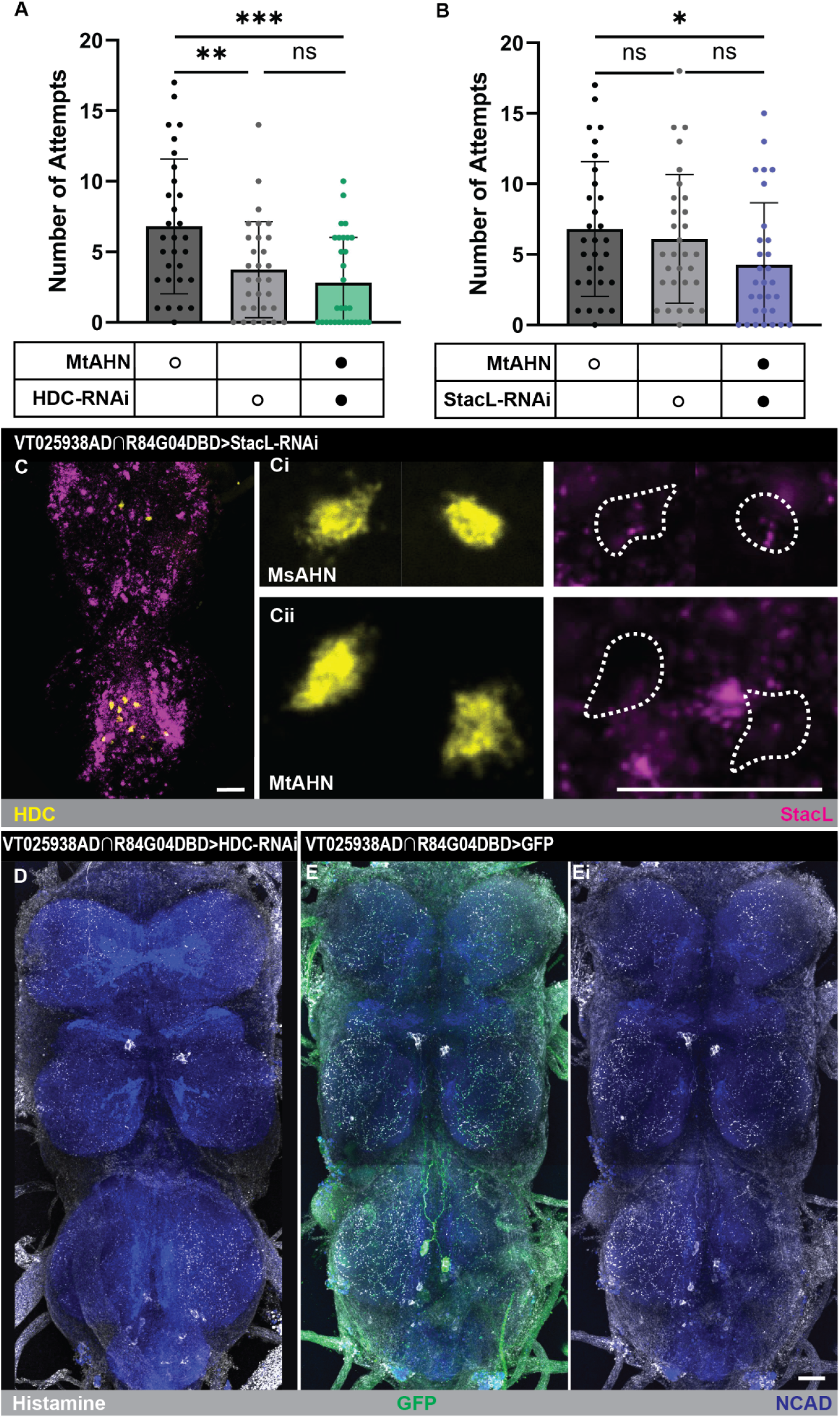
Number of courtship attempts does not change with RNAi mediated knock down of HDC and StacL. IHC and HCR labeling confirms HDC and StacL-RNAi knock down expression in MtAHNs. (A-B) Bar graphs of number of courtship attempts per fly separated by genotype (below). **(A)** Comparison of courtship attempts between VT025938AD∩R84G04DBD driving HDC-RNAi and Gal4 and UAS controls. **(B)** Comparison of courtship attempts between VT025938AD∩R84G04DBD driving StacL-RNAi and Gal4 and UAS controls. Significance from t-tests indicated above (N=30, significance from one-way ANOVA with post-hoc Tukey’s test for individual comparisons: NS=P>0.05, *=P<0.05, **=P<0.01, ***=P<0.001) **(C)** VNC image of HCR labeling for HDC (yellow) and StacL (magenta) in VT025938AD∩R84G04DBD driving StacL-RNAi expression with cell body images of the **(Ci)** MsAHNs and **(Cii)** MtAHNs, cell body locations indicated with white dotted lines. **(D)** Histamine antibody labeling (white) in the VNC of VT025938AD∩R84G04DBD driving HDC-RNAi expression. **(E-Ei)** VNC image of VT025938AD∩R84G04DBD driving GFP expression (green) with antibody labeling for Histamine (white) and NCAD (Blue) as a neuropil marker. Scale bars=20μm.

**Figure S6.1.**
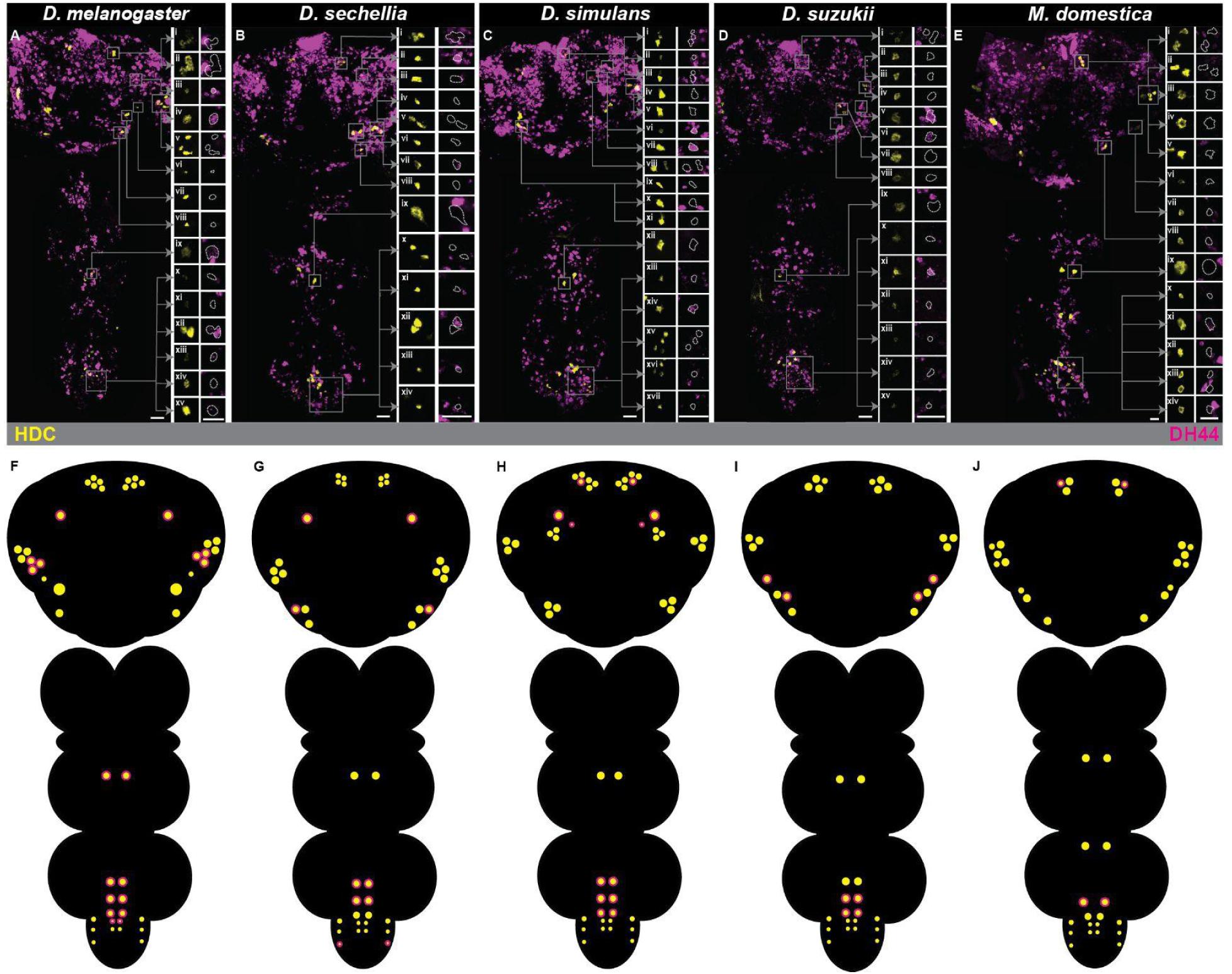
Overlap of HDC and Dh44 in the full CNS of species of *Drosophila* and *M. domestica*. (A-E) Full CNS HDC (yellow) and Dh44 (magenta) HCR labeling with species indicated above. **(Ai-viii, Bi-viii, Ci-xi, Di-viii, Ei-viii)** Cell bodies of the HDC+ (yellow) cells in the brain with Dh44 (magenta) HCR labeling. HDC cell location indicated with white dotted lines. **(Aix, Bix, Cxii, Dix, Eix)** Cell bodies of the MsAHNs with HDC (yellow) and Dh44 (magenta) HCR labeling. HDC cell location indicated with white dotted lines. **(Ax-xv, Bx-xiv, Cxiii-xvii, Dx-xv, Ex-xiv)** Cell bodies of the abdominal HDC+ cells (yellow) with Dh44 (magenta, HDC cell location indicated with white dotted lines) HCR labeling. Cell body scan location indicated with grey boxes. Scale bars = 10μm. **(F-J)** Schematics of the CNS of the species above with HDC+ cells indicated with yellow circles and Dh44 overlap indicated with magenta outline.

**Figure S6.2.**
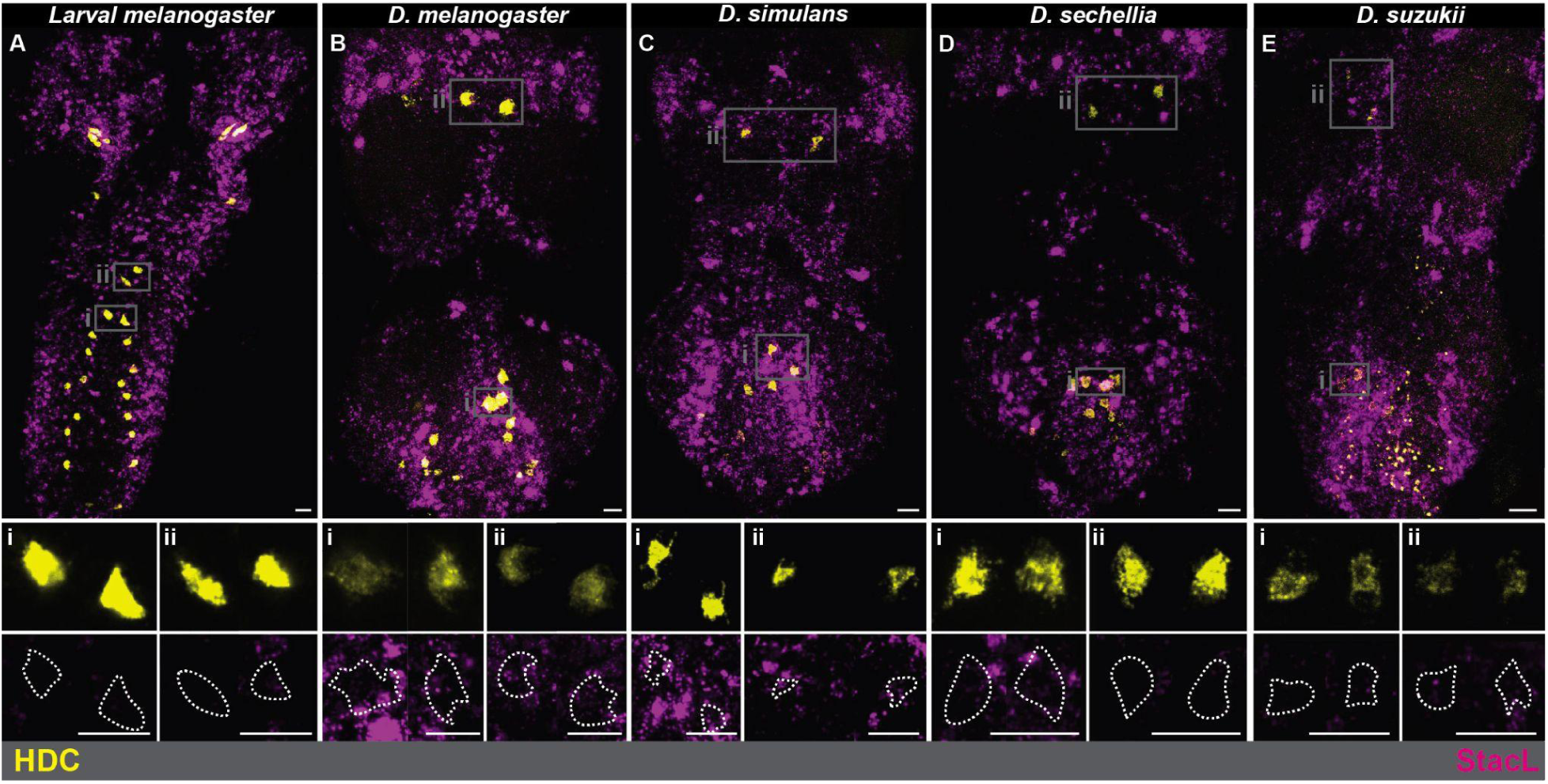
StacL expression in the larval AHNs and other Drosophila species. (A-E) HDC (yellow) and StacL (magenta) labeling in the VNC in the species indicated above. **(Ai-Ei)** MtAHN and **(Aii-Eii)** MsAHN cell body images with HDC (yellow) and StacL (magenta) HCR labeling. Higher magnification scan location indicated with grey boxes and HDC labeling of MtAHNs indicated with white dotted line. Scale bars = 10μm.

## Supplementary Tables

**Table S1.** Top downstream partners (>20 synapses) of the MtAHNs in MANC.

| ID | MtAHN Synapses | MANC type | Cell Class | Instance |
| --- | --- | --- | --- | --- |
| 10098 | 65 | AN02A001 | AN | AN02A001_T2_L |
| 10137 | 59 | AN02A001 | AN | AN02A001_T2_R |
| 10097 | 40 | DNxl080 | DN | DNxl080_CvC_L |
| 10757 | 35 | IN17B003 | IN | IN17B003_A2_L |
| 10606 | 30 | IN17B003 | IN | IN17B003_A2_R |
| 11038 | 29 | IN21A010 | IN | IN21A010_T2_L |
| 26633 | 28 | IN13B080 | IN | IN13B080_T2_R |
| 11122 | 28 | IN21A004 | IN | IN21A004_T3_R |
| 10978 | 27 | IN21A004 | IN | IN21A004_T3_L |
| 10117 | 27 | DNxl080 | DN | DNxl080_CvC_R |
| 10054 | 26 | IN17B001 | IN | IN17B001_A2_L |
| 10141 | 25 | IN12B002 | IN | IN12B002_T2_R |
| 101794 | 24 | IN19A011 | IN | IN19A011_T1_L |
| 11143 | 24 | IN04B002 | IN | IN04B002_T1_L |
| 10223 | 24 | AN17B002 | AN | AN17B002_A1_L |
| 10477 | 24 | IN00A001 | IN | IN00A001_T2_M |
| 14095 | 24 | MNad34 | MN | MNad34_AbN2_R |
| 12235 | 23 | SNpp30 | SN | SNpp30_ADMN_L |
| 10532 | 23 | DNg74 | DN | DNxl040_CvC_L |
| 11081 | 23 | IN04B002 | IN | IN04B002_T1_R |
| 11171 | 22 | IN21A004 | IN | IN21A004_T2_L |
| 10642 | 21 | INXXX038 | IN | INXXX038_A2_L |
| 10187 | 21 | IN21A004 | IN | IN21A004_T2_R |

**Table S2.**
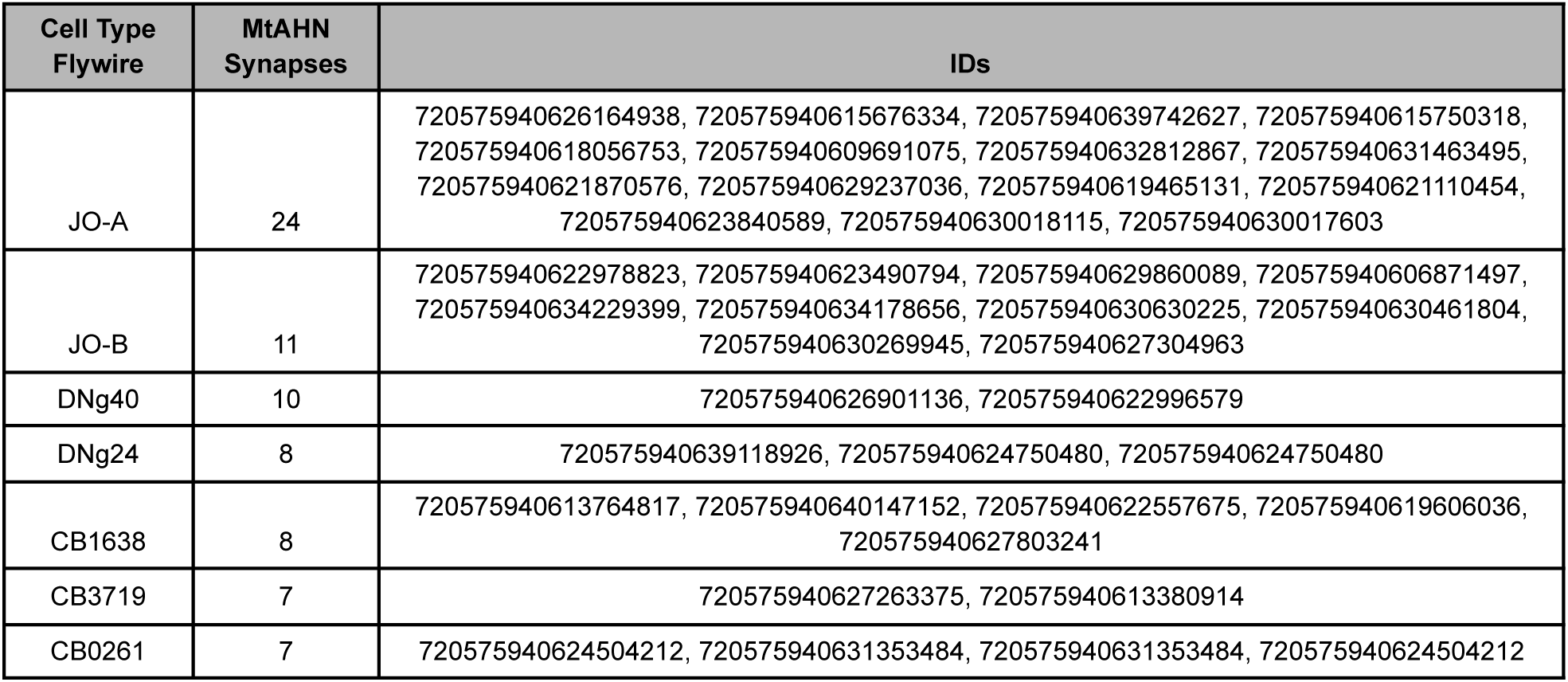
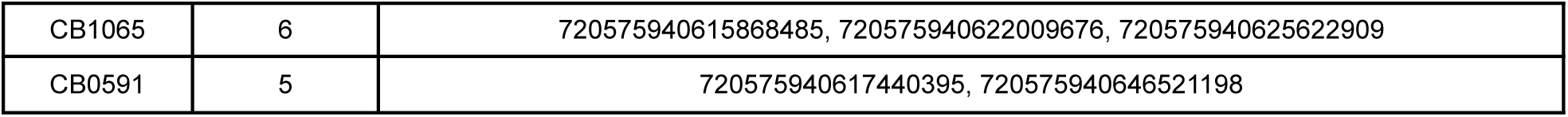
Top cell types downstream of the MtAHNs in the AMMC in FAFB.

**Table S3.**
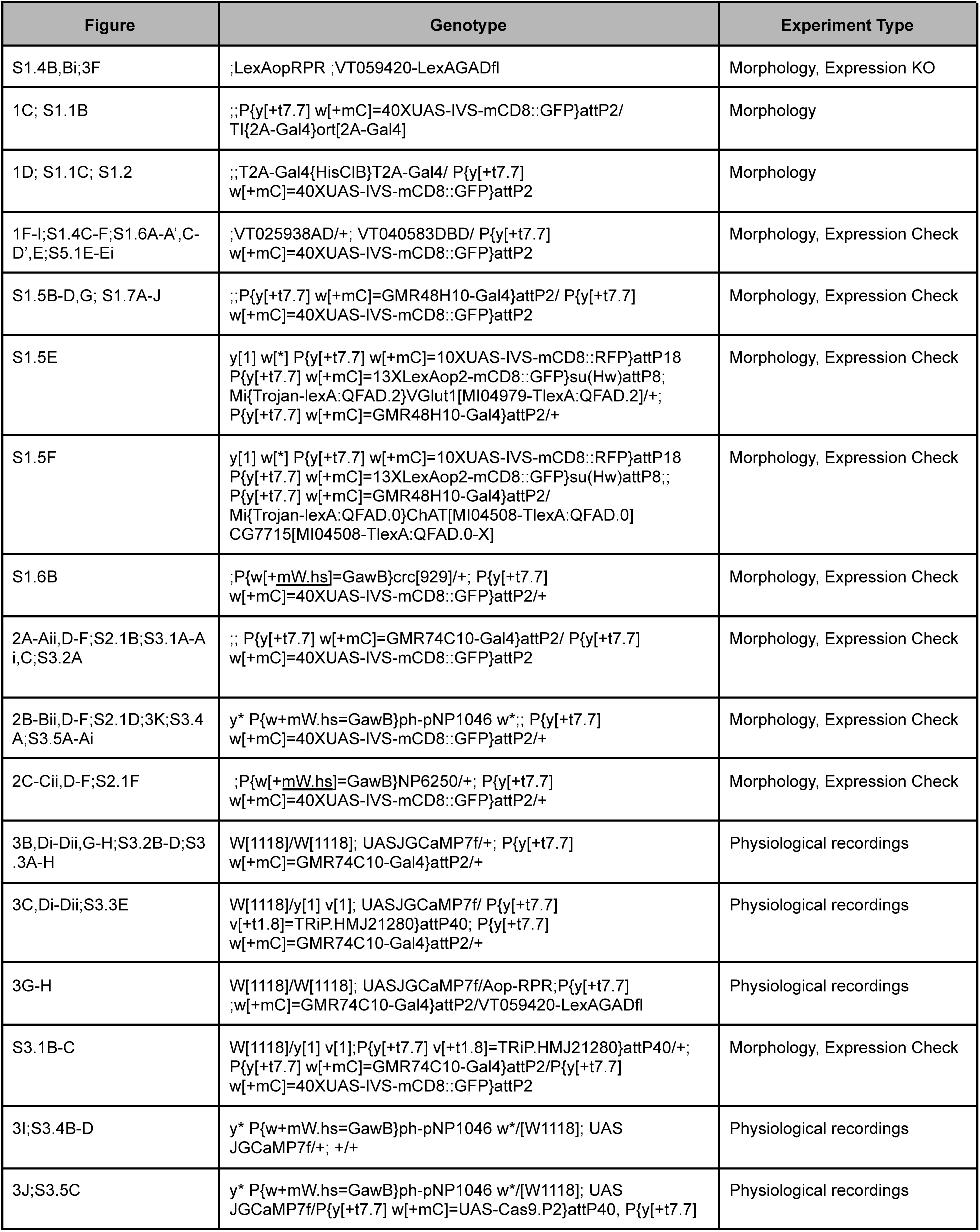
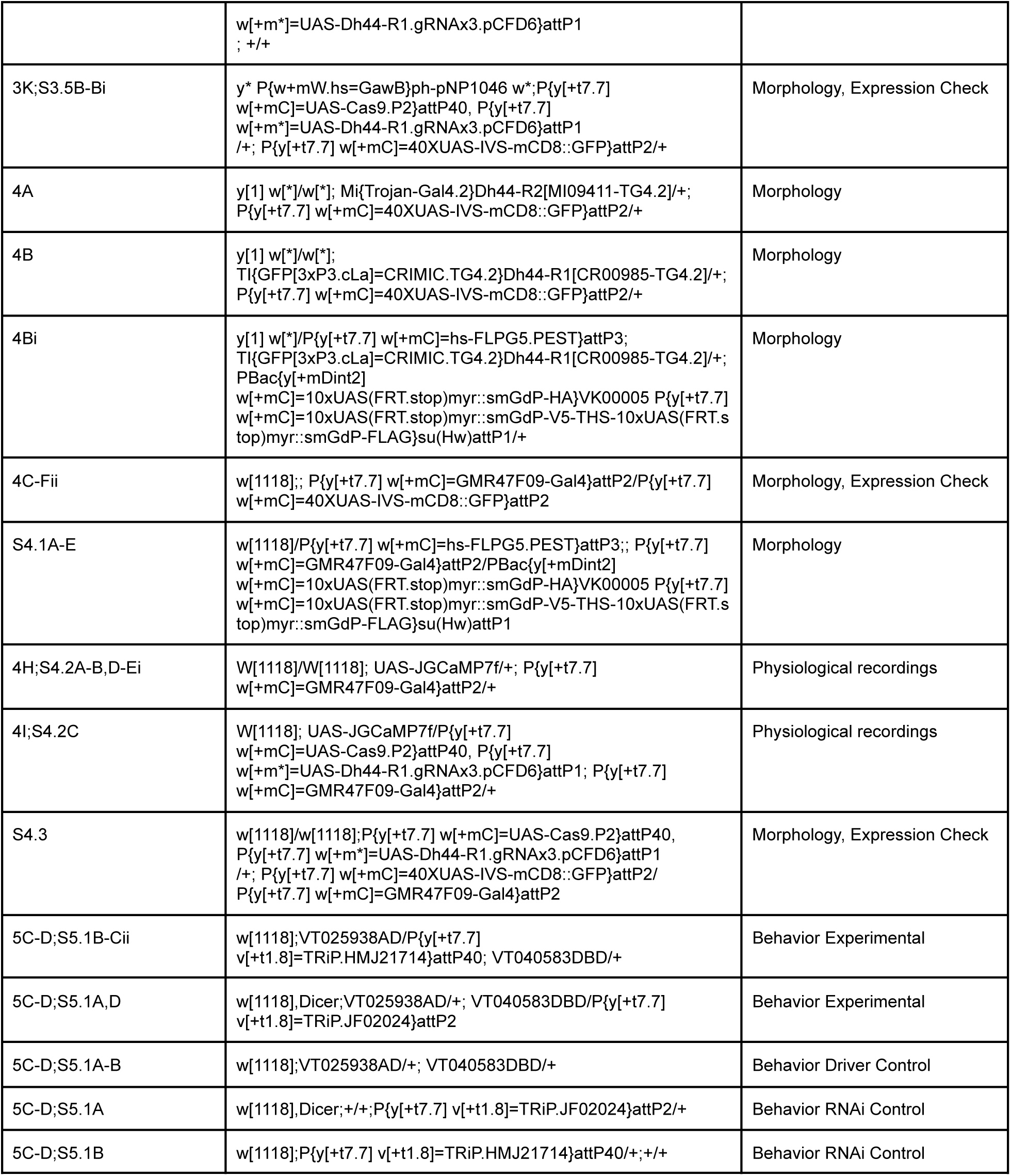
Genotypes per figure.

